# Binary interactome models of inner- versus outer-complexome organisation

**DOI:** 10.1101/2021.03.16.435663

**Authors:** Luke Lambourne, Anupama Yadav, Yang Wang, Alice Desbuleux, Dae-Kyum Kim, Tiziana Cafarelli, Carles Pons, István A. Kovács, Noor Jailkhani, Sadie Schlabach, David De Ridder, Katja Luck, Wenting Bian, Yun Shen, Zhipeng Yang, Miles W. Mee, Mohamed Helmy, Yves Jacob, Irma Lemmens, Thomas Rolland, Atina G. Coté, Marinella Gebbia, Nishka Kishore, Jennifer J. Knapp, Joseph C. Mellor, Jüri Reimand, Jan Tavernier, Michael E. Cusick, Pascal Falter-Braun, Kerstin Spirohn, Quan Zhong, Patrick Aloy, Tong Hao, Benoit Charloteaux, Frederick P. Roth, David E. Hill, Michael A. Calderwood, Jean-Claude Twizere, Marc Vidal

## Abstract

Hundreds of different protein complexes that perform important functions across all cellular processes, collectively comprising the “complexome” of an organism, have been identified^1^. However, less is known about the fraction of the interactome that exists outside the complexome, in the “outer-complexome”. To investigate features of “inner”- versus outer-complexome organisation in yeast, we generated a high-quality atlas of binary protein-protein interactions (PPIs), combining three previous maps^2–4^ and a new reference all-by-all binary interactome map. A greater proportion of interactions in our map are in the outer-complexome, in comparison to those found by affinity purification followed by mass spectrometry^5–7^ or in literature curated datasets^8–11^. In addition, recent advances in deep learning predictions of PPI structures^12^ mirror the existing experimentally resolved structures in being largely focused on the inner complexome and missing most interactions in the outer-complexome. Our new PPI network suggests that the outer-complexome contains considerably more PPIs than the inner-complexome, and integration with functional similarity networks^13–15^ reveals that interactions in the inner-complexome are highly detectable and correspond to pairs of proteins with high functional similarity, while proteins connected by more transient, harder-to-detect interactions in the outer-complexome, exhibit higher functional heterogeneity.

---

Intracellular organisation relies on large numbers of interactions amongst protein, RNA, and DNA macromolecules, forming complex networks that underlie diverse functional relationships. Efforts to map biophysical interactome networks, such as protein-protein interaction (PPI) networks^16, 17^ helped advance our understanding of cellular organisational principles^18, 19^. However, most current models of PPI networks ignore the wide range of biophysical properties exhibited by PPIs, of which the principal dimension is the difference between stable complexes and transient interactions. The interactome is often conceived of as a collection of hundreds of multimeric machines, collectively referred to as the “complexome”^1^. However, stable PPIs forming quaternary structures are only a subset of the protein interactome^20, 21^. There are also weaker transient interactions that are much more dependent on the cellular context, and that are more likely to underlie the overall organisation and compartmentalization in cells and help sustain biochemical pathways and signalling cascades^22–24^.

It has been shown that transient interactions outnumber interactions in stable complexes in the human interactome^25^. However, little is currently known about how these two categories of PPIs underlie different functional relationships between proteins. To address this, we selected *S. cerevisiae*, which has the most comprehensive and diverse systematic datasets of functional relationships between genes as well as multiple large-scale maps of PPIs. We compare intra-complex interactions taking place “inside” each complex of the complexome, i.e. within the “inner-complexome”, to interactions between proteins in different complexes, or involving proteins not known to be in any complex, i.e. within the “outer-complexome”. To facilitate this, we generated for the first time an “all-versus-all” systematic binary interactome map. By integrating this, and other, PPI networks with global functional similarity networks, we found strong support for an emerging model^25–28^ in which a relatively small proportion of the interactome corresponds to the inner-complexome, connecting functionally homogenous proteins, while the vast majority of the interactome consists of interactions between functionally heterogeneous proteins in the outer-complexome.

## The inner- and outer-complexome

It is thought that most cellular processes are carried out by multiprotein molecular machines^6^. However, across different species, the large majority of proteins have not been shown to participate in any protein complex comprising three or more proteins, using a manually-curated dataset of protein complexes (Fig. 1a). After combining multiple protein complex datasets, to reduce the false negative rate as much as possible, proteins outside of complexes are still in the majority (Extended Data Fig. 1a). This has major implications when considering how PPI networks underlie cellular function, as it suggests a large fraction of the functions of interactions are performed outside complexes. To categorise PPIs in relation to protein complexes, we divide all pairs of proteins into four different “zones” (Fig. 1b): Zone A, the inner-complexome, corresponds to all pairwise combinations where both proteins are subunits of the same complex; Zone B corresponds to pairs of complex subunits where each protein belongs to a different complex; Zone C represents all pairwise combinations involving both a complex subunit and a non-complex protein; Zone D corresponds to all pairwise combinations between non-complex proteins. In yeast, the inner-complexome, Zone A, corresponds to ∼17,600 protein pairs, representing 0.1% of the approximately 18 million possible protein pairs, with the remaining 99.9% of pairs residing in the outer-complexome – Zones B, C, and D (Fig. 1b) – highlighting the enormous potential role that outer-complexome interactions play in eukaryotic biology.

**Fig. 1.**
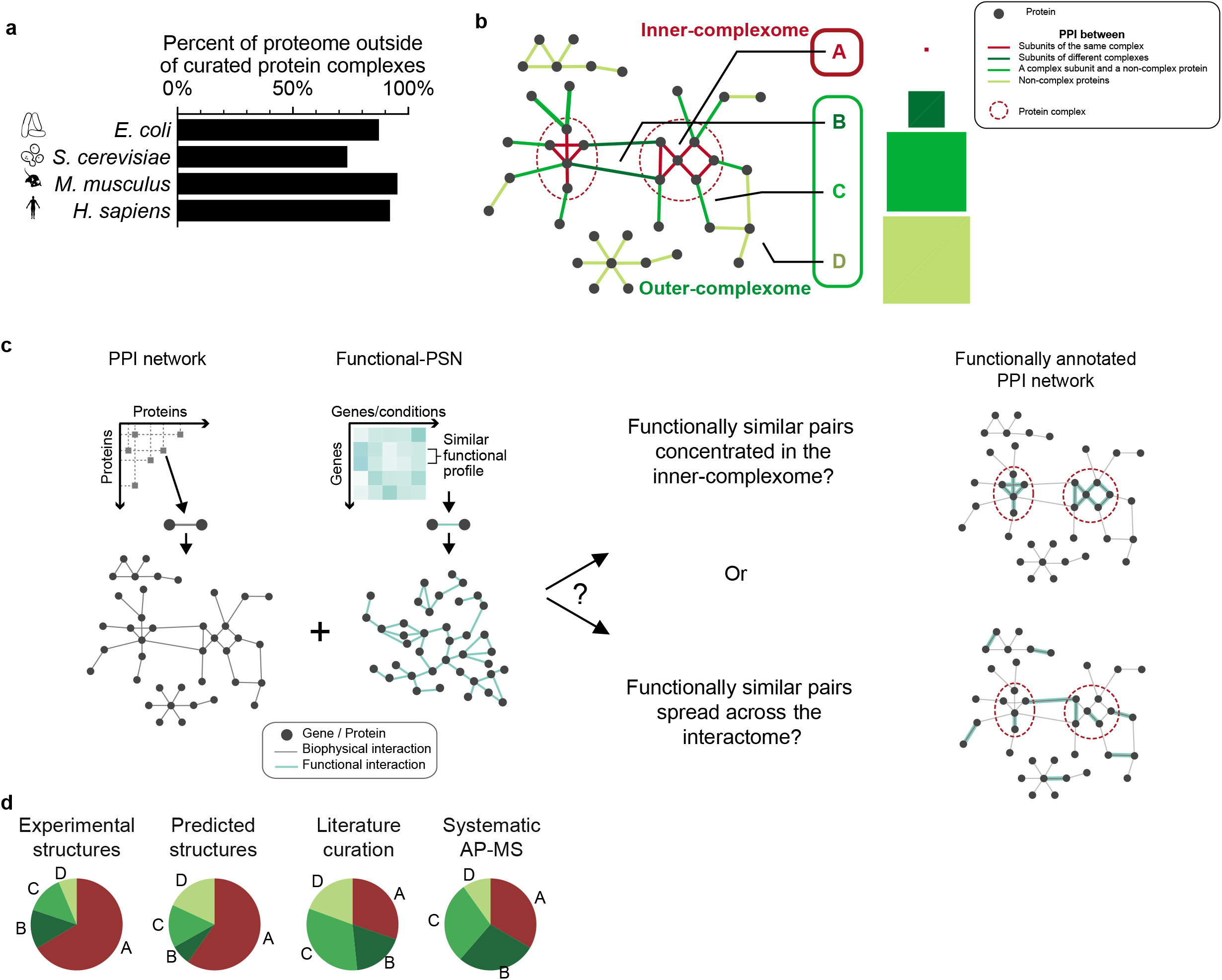
The inner- and outer-complexome. **a**, The fraction of the proteome of different organisms that are not listed as subunits of protein complexes of at least 3 or more different protein subunits. **b,** Illustrative network diagram showing the space of pairwise protein combinations categorised into four Zones based on protein complex membership. The area of each square is proportional to the size of the number of pairwise combinations of proteins in each Zone, for yeast. **c**, Schematic of the approach and question of functional similarity of interacting proteins by combining biophysical and functional networks. **d**, The proportion of four different yeast biophysical interaction datasets in each Zone.

We investigate the roles of inner- and outer-complexome PPIs in the interactome by, first, understanding the distribution of biophysical PPIs in the different zones, through a detailed assessment of available PPI networks. Next, we examine whether there is a difference in the functional relationships between physically interacting proteins in the different zones, using functional profile-similarity networks (PSNs) (Fig 1c). We considered four different sources: i) experimentally resolved 3D structures^29^ (I3D-exp-20); ii) recent deep learning-based prediction of PPI structures^12^ (Alphafold+RoseTTAFold); iii) literature-curated binary pairs supported by multiple pieces of evidence^8–11, 30^ (Lit-BM-20); and iv) systematic large scale AP-MS (Sys-NB-06) (Supplementary Tables 1-5). These datasets show variation in their distribution between these four zones, with around one-third of pairs in Lit-BM-20 and Sys-NB-06 are between subunits of the same complex (Zone A), compared to more than half in I3D-exp-20 and AlphaFold+RoseTTAFold, (Fig. 1d).

## All-by-all yeast reference interactome

These results present conflicting views of what fraction of a eukaryotic interactome exists outside the inner-complexome. To resolve these differences, we turned to another type of dataset: systematic binary PPI maps, generated using yeast two-hybrid (Y2H) as the main screening assay. We hypothesise that uniform testing of the entire proteome-by-proteome space should give the most unbiased estimate of the proportion of the interactome in different zones. However, the three systematic Y2H maps currently available^2–4^, collectively referred to as Y2H-union, were obtained using incomplete sets of open reading frames (ORFs), or “ORFeome” collections^31^, each screening only ∼70-75% of the search space. To systematically generate a map that could provide a complete and unbiased estimate of the proportion of the interactome in different zones, we started by compiling a high-quality ORFeome collection covering 99% of yeast protein-coding genes, by verifying an existing collection of 4,933 ORFs^32^, and cloning an additional 921 ORFs (Fig. 2a, Supplementary Table 6). To maximize the potential for novel discovery relative to the three previous binary systematic maps, we implemented a new assay version, Y2H v4, and demonstrated that it detected an orthogonal set of interactions compared to previous Y2H versions (Extended Data Fig. 2a, Supplementary Table 7). We systematically screened 27 million bait-prey combinations three times independently. Pairs identified in these primary screens were subsequently evaluated in two independent pair-wise tests. To ensure a high-quality reproducible dataset, only pairs that scored positive and were sequence-confirmed in both attempts were considered positive. The quality of the dataset was assessed by testing every pair in two orthogonal binary PPI assays – MAPPIT^33^ and GPCA^34^ – alongside positive and random reference sets (Fig. 2b, c, Extended Data Fig. 2b, Supplementary Tables 8-10). Thus, we generated a new yeast interactome map of 1,910 PPIs between 1,351 proteins, which we term the Yeast Reference Interactome (YeRI) (Supplementary Table 11). Three-quarters of the PPIs in YeRI are novel. YeRI pairs tested positive in orthogonal assays at rates similar to Lit-BM pairs available in 2013 (Lit-BM-13). YeRI displays significant enrichment for interactions between proteins that share annotations for cellular compartments, pathways, or protein complexes (Extended Data Fig. 2c), demonstrating its overall high level of biological relevance.

**Fig. 2.**
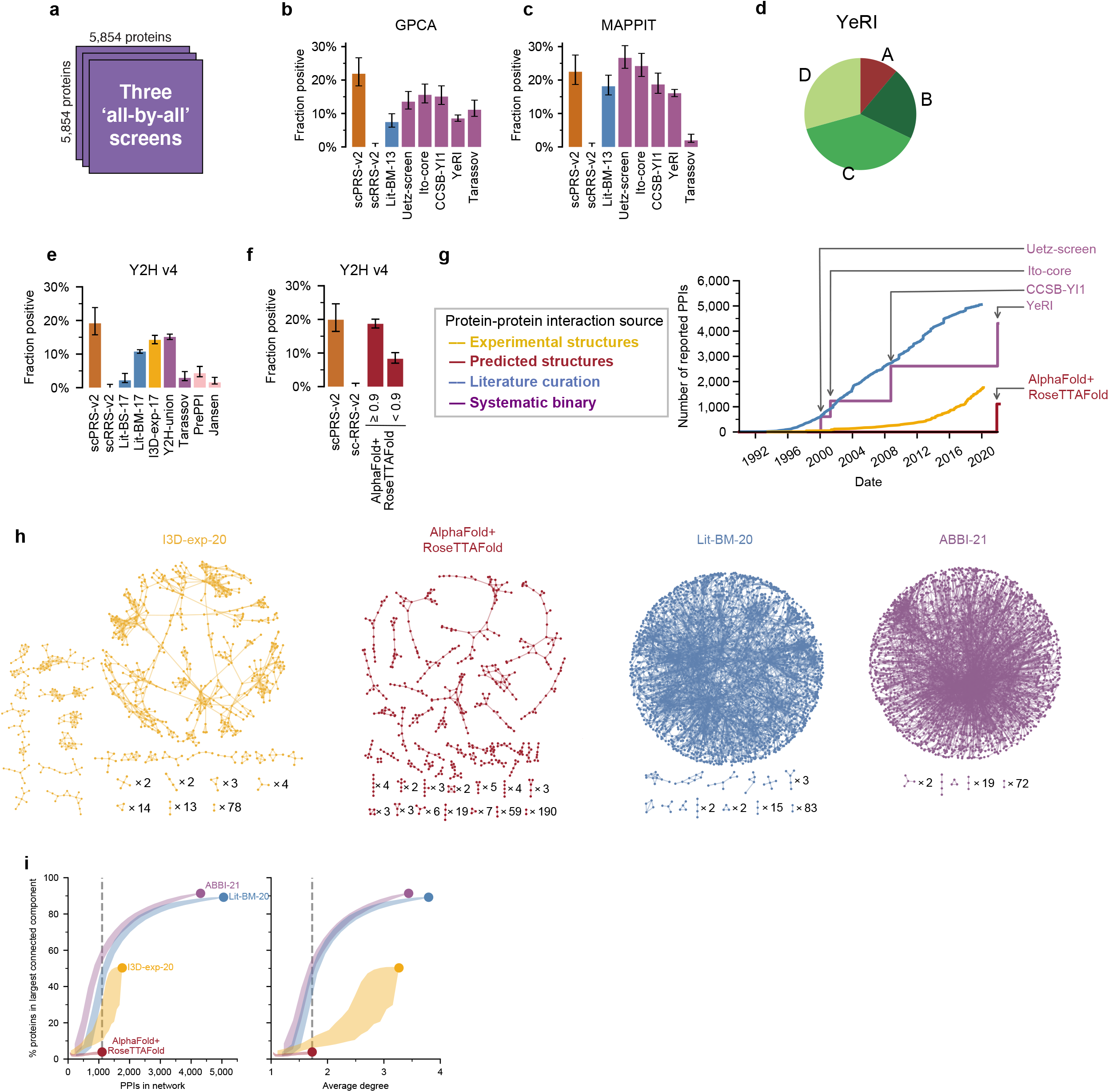
All-by-all yeast reference interactome and comparison to existing high-quality binary interactome maps. **a**, The YeRI screening space. **b**, **c**, Experimental validation of yeast systematic binary maps and literature-curated pairs in GPCA (**b**) and MAPPIT (**c**). Error bars are 68.3% Bayesian credible intervals. **d**, The proportion of YeRI in each Zone. **e**, Yeast PPI datasets tested in Y2H v4. Error bars are 68.3% Bayesian credible intervals. **f**, Results of testing predicted structure dataset in Y2H v4. Error bars are 68.3% Bayesian credible intervals. **g**, Increase in the number of high-quality binary PPIs from available experiments in Interactome-3D (yellow), AlphaFold+RoseTTAFold (brown), Lit-BM (blue), and systematic studies (purple). **h**, Networks for four different high-quality binary PPI datasets. **i**, Fraction of proteins in the largest connected component, with points showing the values for four PPI networks, and shaded bands showing random subsamples of each network across a range of sizes, plotted against the number of PPIs (left) and the average degree per protein (right). Shaded bands indicate the innermost 95% interval of random subnetworks.

Since we showed that all four systematic Y2H maps are well validated by MAPPIT and GPCA (Fig. 2b, c), we combined them into a single Atlas of Binary Biophysical Interactions (ABBI-21) comprising 4,556 PPIs (Supplementary Table 12). Clusters of interacting proteins enriched for shared Gene Ontology (GO) terms^35^, increase substantially with the addition of each map (Extended Data Fig. 2d, e). Due to the expanded set of screened ORFs, YeRI substantially improves coverage for genes of unknown function and can be used to predict their functions (Extended Data Fig. 2f-h, Supplementary Note 1, Supplementary Table 13). In total, ABBI-21 covers 12-25% of the estimated yeast binary interactome (Extended Data Fig. 3a)^4, 36, 37^. The proportion of pairs in different zones from YeRI and ABBI-21 (Fig. 2d, Extended Data Fig. 3b) produces a substantially different picture of the interactome compared to the other maps (Fig. 1d), with only around 10% of PPIs in Zone A and substantially more in Zones C and D than the other networks. This view suggests that the outer-complexome is dominant in terms of the number of PPIs in the interactome.

**Fig. 3.**
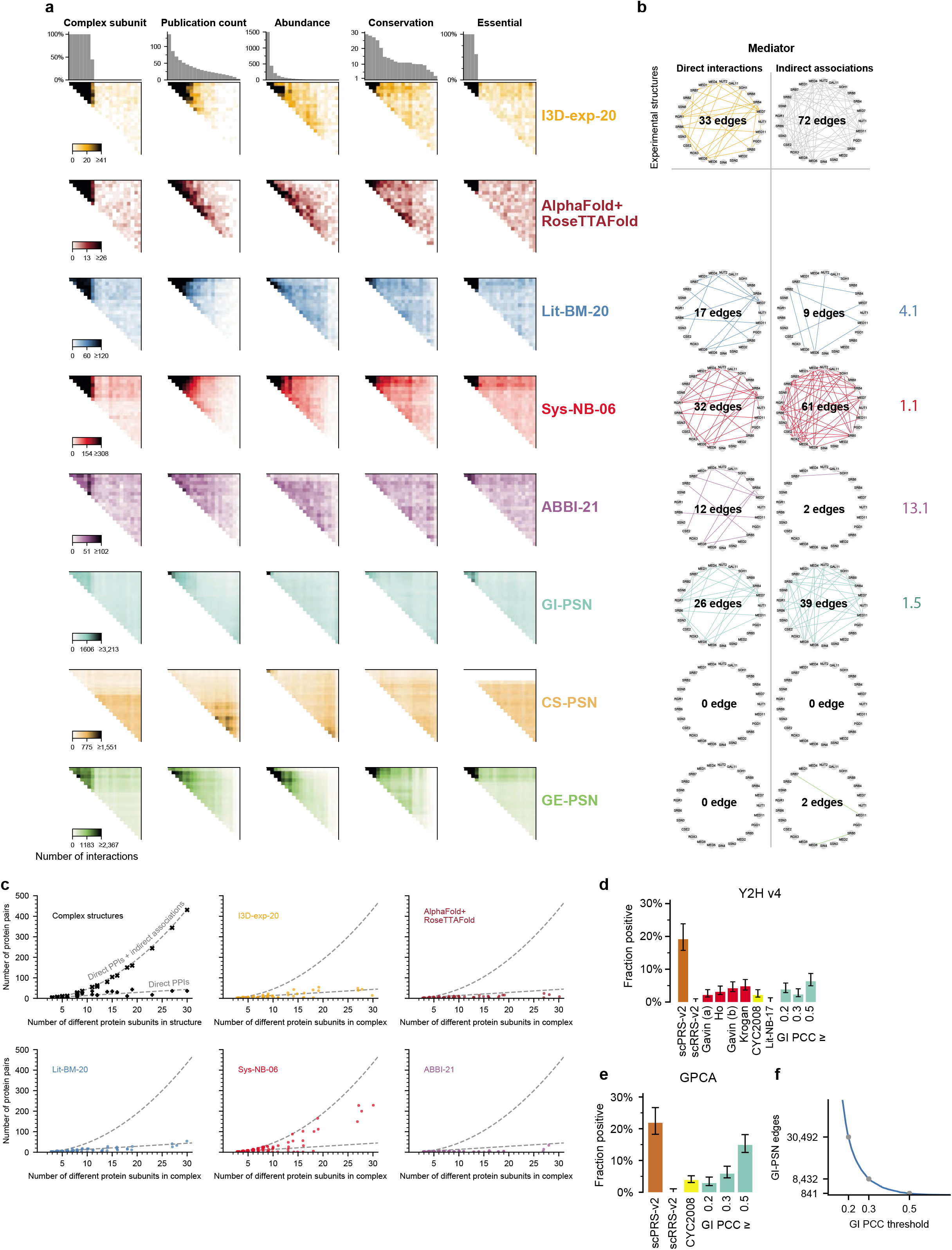
High discrepancy among alternative views of interactome organisation. **a**, Heatmaps of protein interactions and associations in biophysical and functional networks, ordered by complex membership, the number of publications per protein, protein abundance, gene conservation, and essentiality. Upper limits of the colour scales are set to 5x the average number of PPIs per bin, separately for each network. **b,** Using experimental structures of mediator, restricting to pairs of proteins in the same structure, the direct interactions and indirect associations (top row) and of those, which ones are contained within the different biophysical and functional networks (rows below). Direct enrichment is defined as the ratio of the fraction of detected direct interactions over the fraction of detected indirect associations. **c**, Top-left panel: the number of either direct (+) or both direct and indirect (x) PPIs in the single largest experimental structure for each protein complex. The curved dashed line corresponds to the total possible pairwise combinations of different proteins per complex; the straight dashed line is a linear regression using the direct PPIs. Remaining panels: the number of PPIs in each of the five networks within each complex, against complex size. The dashed lines from the top-left panel are reproduced. **d**, **e**, Results of testing samples of pairs from GI-PSNs, across different PCC cutoffs, in Y2H v4 (**d**) and GPCA (**e**). Error bars are 68.3% Bayesian credible intervals. **f**, The number of edges in GI-PSN at the tested PCC cutoffs.

In addition to ABBI-21, to ensure our conclusions would be robust, we also wanted to use alternative high-quality binary PPI datasets in our investigation of the differences of the interactome between the inner- and outer-complexome. So we performed a comprehensive test of the biophysical quality of available sources of binary PPI data. We experimentally retested all pairs available in 2017 for I3D-exp, Lit-BM and Y2H-union, along with random samples of other datasets, testing a total of 8,999 pairs in Y2H v4 (Fig. 2e, Extended Data Fig. 3c, Supplementary Table 14). Y2H v4 recovered 14%, 11%, and 15% of pairs from experimental structures, literature curation, and systematic binary pairs, respectively. These recovery rates are quite high, approaching assay sensitivity limits: indeed, the systematic Y2H datasets were indistinguishable from both scPRS-v2 (*P* = 0.2, one-sided Fisher’s exact test) and I3D-exp-17 (*P* = 0.6, two-sided Fisher’s exact test), our benchmarks of true direct, heterodimeric binary interactions, and all three performed slightly better than Lit-BM-17 (*P* = 0.006, two-sided Fisher’s exact test) (Fig. 2e), again demonstrating that the biophysical quality of systematic binary interaction maps is at least as good, if not superior to that of literature-curated binary interactions. Interestingly, a proteome-scale dataset generated using a dihydrofolate reductase protein complementation assay (Tarassov) that detects physically-proximal, but not necessarily directly-contacting protein pairs^38^, was not significantly above the scRRS-v2 negative control in MAPPIT (*P* = 0.12, one-sided Fisher’s exact test) but validated *on par* with the Y2H-union datasets in GPCA and Y2H v4 (Fig. 2b, c, e). As shown previously for human PPIs^39^, a sample of the putative yeast binary PPIs supported by only a single piece of evidence in the literature in (Lit-BS-17) was recovered at a low rate, not statistically different from scRRS-v2 (*P* = 0.1, one-sided Fisher’s exact test), of only 2% (Fig. 2e). Finally, two datasets of predicted PPIs, PrePPI and Jansen et al^40, 41^, tested positive at low levels of 4% and 2%, respectively (Fig. 2e). Based on these results, out of the reported interaction sets we tested, we restrict the binary PPI maps in analysis of the inner- and outer-complexome, to I3D-exp-20, Lit-BM-20, and ABBI-21.

Testing a comprehensive set of PPIs with experimental structures with Y2H v4 allowed us to investigate to what extent structural features affect the sensitivity of the assay to different PPIs. We first saw that, although the number of subunits involved in forming large protein complexes appears to have some impact on the rate of interaction recovery by Y2H v4 (Extended Data Fig. 3d), binary assays can readily detect pairs of interacting proteins even in large complexes. Second, although our dataset was generated with full-length yeast proteins, the detection rate appeared unaffected by whether the structures of interacting proteins had been solved with full-length proteins or fragments (*P* = 0.14, two-sided Kolmogorov-Smirnov test) (Extended Data Fig. 3e). Third, although we observed a trend towards larger interaction interfaces among Y2H v4 positives compared to negatives (*P* = 2 x 10^-6^, two-sided Mann-Whitney U test), Y2H v4 could detect interactions with interface areas ranging widely from 100 to 10,000 Å^2^ (Extended Data Fig. 3f). Y2H v4 was better able to detect PPIs with small interaction interfaces than GPCA (*P* = 0.0043, two-sided Mann-Whitney U test) and was at least as good at this as MAPPIT (*P* = 0.11, two-sided Mann-Whitney U test). Finally, Y2H v4 detected interactions with *K_d_* values up into the micromolar range (Extended Data Fig. 3g), consistent with previous findings that Y2H can identify weak interactions^7, 42^. Together, the high and consistent sensitivity of Y2H v4 to a variety of different PPIs, across strong and weak interactions, and within and outside protein complexes, suggest that YeRI is representative of the real interactome across both the inner- and outer-complexome.

Beyond testing PPIs with experimental structures, we next used Y2H v4 to perform the first experimental assessment of the quality of a recent deep learning-based dataset of PPIs with predicted structures^12^. Complex structure prediction represents a new and promising method for generating detailed PPI data, so it is crucial to have a good understanding of its quality. The AlphaFold+RoseTTAFold predicted structures have a contact probability for each pair, and we observed that pairs with a contact probability above 0.9 test positive at a rate in line with scPRS-v2 (Fig. 2f, Extended Data Fig. 3h, Supplementary Table 15). Therefore we restricted subsequent analysis of AlphaFold+RoseTTAFold to the 1,106 out of 1,505 PPIs (73%) with contact probability ≥ 0.9, of which 392 PPIs (35%) have no previous experimental structure available (Extended Data Fig. 3i). Comparing the resulting AlphaFold+RoseTTAFold network to I3D-exp-20, Lit-BM-20, and ABBI-21, we observed that the structural data, both experimental and predicted, produce more fragmented networks, largely due to a lack of highly connected hub proteins in those networks (Fig. 2g-i, Extended Data Fig. 3j).

## Alternative views of interactome organisation

To correctly infer the distribution of PPIs between the inner- and outer-complexome, we need to understand any potential biases in the coverage of the different networks. It has been shown for human PPIs that networks generated by varying methods can be concentrated within a subset of the space of pairwise protein combinations^39^, but this has not yet been investigated in yeast. So we displayed binary PPIs in a representation of the proteome-by-proteome space that was organised by ranking proteins in both dimensions according to complex membership and four protein properties: number of publications, abundance, evolutionary conservation, and essentiality (Fig. 3a, Supplementary Table 16). Complex subunits exhibit significantly higher average values in all four variables (*P* < 10^-83^, Mann-Whitney *U* test), making them more well-studied, more abundant, more conserved, and more often essential than non-complex proteins (Extended Data Fig. 4a). There are strong correlations between these protein properties (Extended Data Fig. 4b). When ranking by publications per gene, all datasets show a much higher density of PPIs between highly-studied proteins, except ABBI-21, which is distributed more homogeneously. To quantify this, the size of the matrix of the most studied proteins containing 80% of PPIs from each dataset is 16%, 37%, 25%, 25%, and 57% for I3D-exp-20, AlphaFold+RoseTTAFold, Lit-BM-20, Sys-NB-06, and ABBI-21, respectively. Although Sys-NB-06 was obtained using sociologically unbiased, systematic approaches, it nevertheless shows a bias towards highly studied proteins, presumably because detection of complexed proteins using MS-based methods is more sensitive for highly expressed proteins, which also tend to be more highly studied^20^. In all cases, ABBI-21 covers the interactome more homogeneously, across multiple biological properties, compared to Lit-BM-20, Sys-NB-06, and AlphaFold+RoseTTAFold. ABBI-21 does show some depletion for the highest abundance proteins as well as the lowest abundance, least studied, and least conserved proteins (Fig. 3a). In the lowest abundance or conservation zones, the number of proteins with at least one interactor is significantly higher for ABBI-21 than Lit-BM-20, Sys-NB-06, or AlphaFold+RoseTTAFold (Extended Data Fig. 4c). Using a different approach, we tested the recovery of four gold-standard curated sets of PPIs from either the inner- or outer-complexome in the systematic biophysical datasets, and we observed that, although both ABBI-21 and Sys-NB-06 capture the inner-complexome pairs more readily than outer-complexome pairs, ABBI-21 shows more uniformity between inner- and outer-complexome pairs than Sys-NB-06 (Extended Data Fig. 4d).

**Fig. 4.**
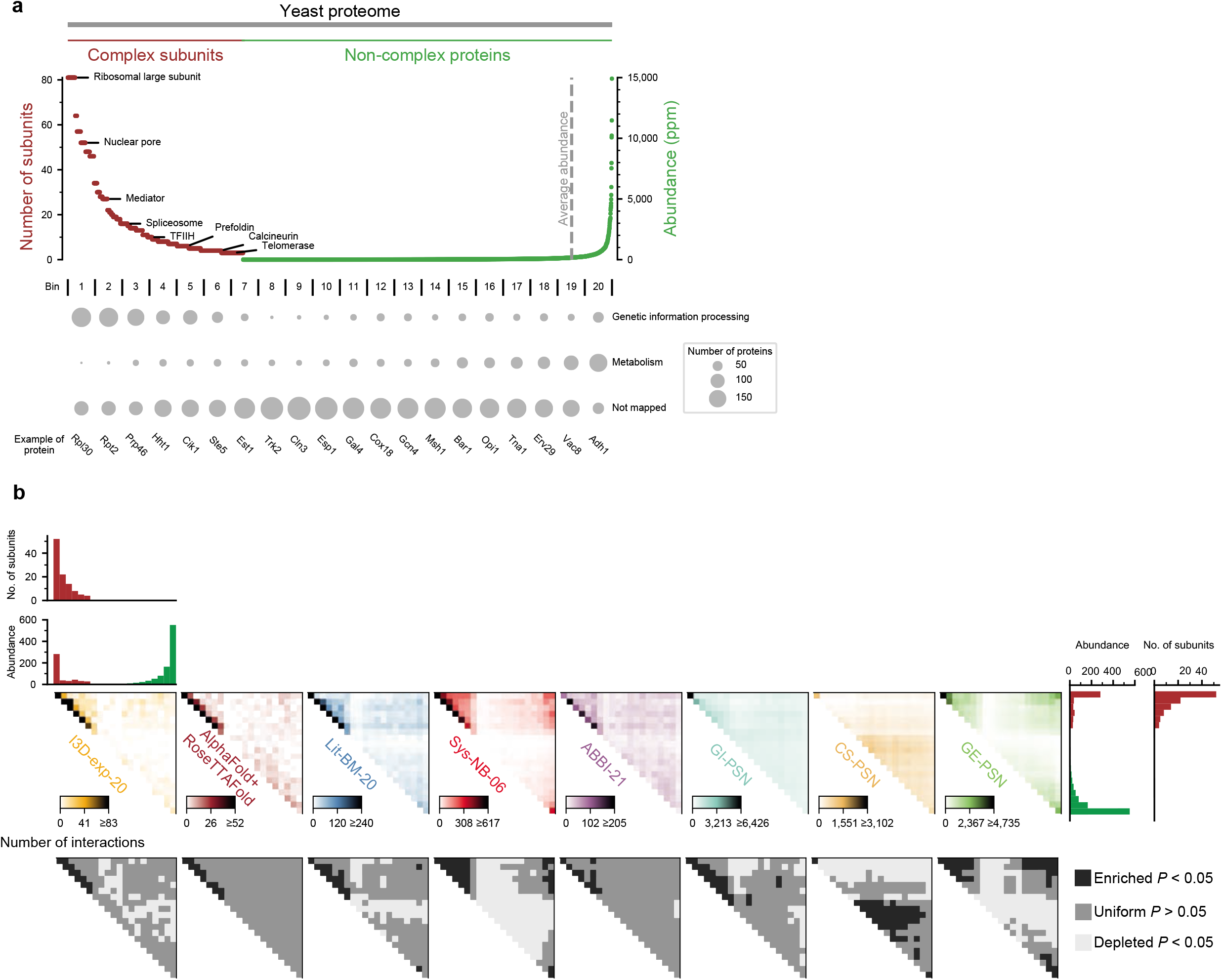
Organisation within the inner- and outer-complexome. **a**, The yeast proteome ordered by decreasing number of subunits of the corresponding complex, for proteins in a complex, and by increasing abundance, for non-complex proteins. After dividing into 20 bins, the number of proteins in each of three general functional categories^62^ is shown by the area of the circles. Not Mapped corresponds to unannotated proteins. **b**, Upper: heatmaps of the number of connected gene pairs in biophysical and functional networks, with the proteome first ordered by size of protein complexes in which the proteins are involved, then ordered by protein abundance. Lower: statistically significant enrichment and depletion of edges per bin, calculated by random permutation of the order of the proteins.

Having identified major differences between the coverage of the proteome-by-proteome space between different PPI networks, we then turned to the functional profile similarity networks (PSNs), to see if there were any biases in their coverage that could impact our interpretation of the functional differences between inner- and outer-complexome PPIs from integrating the PSNs with the PPI networks. The PSNs used three different systematic genome-wide functional genomic profiling approaches, based on: i) positive and negative genetic interactions observed in double mutants bearing knock-out (KO) and/or hypomorphic alleles^13^; ii) growth of KOs of non-essential genes across over 1,000 chemical and environmental stress conditions^14^; and iii) transcriptome-wide measurements of gene expression over thousands of samples^15^. PSNs for genetic interactions, “GI-PSN”^13^; condition sensitivity, “CS-PSN”^14^; and gene expression, “GE-PSN”^15^ were generated from the top one percent of the strongest correlation of tested pairs. While these three PSNs exhibit statistically significant overlaps (P < 0.001, one-sided empirical test, restricted to genes tested in all PSNs), these overlaps are small, with only 4% of edges connected in more than one PSN, showing that different PSNs identify complementary functional relationships (Extended Data Fig. 4e). One of the most striking observations was the dense zone exhibited by GE-PSN within the spaces corresponding to highly abundant or conserved proteins as well as essential genes (Fig. 3a). This similarity between Sys-NB-06 and GE-PSN is likely because both experimental strategies are highly dependent on endogenous gene expression levels. The second observation was that GI-PSN also shows a higher density of functional relationships amongst extremely well-studied and highly abundant proteins (Fig. 3a), concentrated in a smaller area than Sys-NB-06. This was unexpected since GI-PSN was generated systematically, independently of any sociological bias. Upon investigation, we found that this was due to a combination of higher connection density for both essential genes^13^ and highly abundant non-essential ribosomal subunits in the GI-PSN network (Extended Data Fig. 4f). Many yeast ribosomal proteins retained paralogs after the whole-genome duplication event^43^, often rendering them non-essential where paralogs can, at least partially, functionally compensate for one another’s deletions. Lastly, CS-PSN was obtained from a homozygous gene deletion collection that does not include essential genes, which could explain some of the observed patterns.

Having observed that different PPI mapping methods have specific biases in their coverage, we then investigated another important distinction between these methods: the difference between detecting co-complex associations where proteins are in the same complex but not necessarily in direct contact, or detecting “binary” PPIs for which two interaction partners are likely to be in direct contact. It is crucial to distinguish between these two types of protein-protein relationship, as it has major consequences for the resulting protein network, with one example being that large protein complexes contain many more indirect associations than direct contacts. We investigated to what extent co-complex and binary biophysical networks identify direct interactions versus indirect associations using available experimental structures of complexes, and compared this with the functional networks. As an example, the mediator complex shows 33 direct contacts between its 25 subunits, which is considerably less than its 300 possible pairwise combinations (Fig. 3b). This is unsurprising, as the number of other proteins a complex subunit may be in contact with is fundamentally limited by the surface area of the subunit. The number of direct interactions within a protein complex scales roughly linearly with the complex’s size, averaging 3 PPIs with each subunit, whereas the number of indirect associations scales quadratically, resulting in a dramatically increasing difference between the two for larger complexes (Fig. 3c). ABBI-21 and Lit-BM-20 primarily find direct interactions, whereas the AP-MS-based Sys-NB-06 reports both direct binary interactions and indirect co-complex association between proteins (Fig. 3b), in roughly equal proportions. The GI-PSN also finds both direct interactions and indirect associations (Fig. 3b), with a preference towards direct PPIs (*P* = 0.02, two-sided Fisher’s exact test), as has been observed previously^44^. That GI-PSN should provide indirect associations stands to reason, as all the proteins in the complex collectively contribute to a common function, irrespective of whether they are in direct contact or not. We then investigated this trend across all different protein complexes for which a 3D structure is available (Extended Data Fig. 4g) and observed that binary PPI datasets primarily find direct-contact pairs, whereas Sys-NB-06, GI-PSN, and GE-PSN connect both direct-contact and indirect co-complex association pairs, with a tendency towards direct PPIs. Among all six datasets analysed here, ABBI-21 is the most enriched for direct PPIs vs indirect associations (*P* = 0.0002, two-sided Wilcoxon signed-rank test).

After observing that, within protein complexes, AP-MS and functional networks correspond to both direct PPIs and indirect associations, we then sought to experimentally confirm this observation by testing random samples of co-complex association and GI-PSN edges using Y2H v4. Pairs from co-complex association datasets were detected at rates much lower than those from binary PPI datasets, although significantly higher than the negative control scRRS-v2 (median *P* = 0.018, one-sided Fisher’s exact test) (Fig. 3d). This result suggests that a large proportion of co-complex association datasets are indirect associations, both in literature-curated protein complexes as in CYC2008^45^, and AP-MS-derived proteome-scale maps^5–7, 46^. This observation is consistent with the result that protein pairs in PPIs obtained by binary assays are two to five times more likely to be in direct contact than co-complex association pairs, using experimental complex structures with at least three subunits (Extended Data Fig. 4h, Supplementary Table 17). For random samples of GI-PSN pairs, across a range of PCC cutoffs, the GPCA test-positive rate increases proportionally to the PCC threshold, with the Y2H v4 test-positive rate is flatter but consistent with an increase with PCC threshold (Fig. 3d, e). For PCC ≥ 0.2 and 0.3, the test-positive rate of GI-PSN pairs in both assays is similar to that of CYC2008. Taken together, these results are consistent with the conclusions that protein complexes dominate the high-PCC GI-PSN pairs^13^, that GI-PSN pairs correspond to both directly-contacting and indirectly-linked complex subunits (Extended Data Fig. 4g), and that higher average PCC values have an increased correspondence with direct interaction as opposed to indirect association^44^. These results provide direct experimental estimates of the fraction of genetic interaction profile similarity relationships that correspond to binary PPIs (Extended Data Fig. 4i). Importantly, at the point where the direct binary PPI content substantially exceeds that of protein complexes, at PCC ≥ 0.5, the GI-PSN contains only 841 edges (Fig. 3f).

## Organisation of inner- and outer-complexome

A huge variety of protein function exists within both complex subunits and non-complex proteins. We next looked at two variables, one within each category. We chose: (i) the size of each protein complex; and (ii) the abundance of non-complex proteins. Both capture some aspect of the continuum between constitutive cellular machinery and context-specific, adaptive, dynamic processes (Fig 4a). Protein complexes span a range of sizes, from 81 different proteins in the ribosomal large subunit to three in telomerase. The abundance distribution is such that a small number of proteins make up a large fraction of expressed proteins detected by mass spectrometry. Of non-complex proteins, 89% are below the mean molar abundance. Among the most abundant non-complex proteins are metabolic enzymes, which, in yeast cells, make up 30% of proteins by molarity, but represent only 10% of all encoded proteins^47, 48^. Just two proteins, the pyruvate kinase encoded by CDC19 and the plasma membrane proton ATPase pump encoded by PMA1, account for more than 2% of the total number of cellular protein molecules. At the other end of the spectrum, lowly expressed proteins such as Cln3, an important cell-cycle regulator, and Ime1, a master regulator of meiosis, are four orders of magnitude lower in abundance than Cdc19 and Pma1.

To visualize the distribution of the networks in the proteome-by-proteome space, relative to these two variables, we ordered complex subunits by decreasing size of the corresponding complex, and non-complex proteins by increasing abundance. All five biophysical datasets show a strong, statistically significant, enrichment for interactions or associations in the inner-complexome (*P* < 0.05, permutation test, Fig. 4b, Extended Data Fig. 5a, b). We observe the same for GI-PSN and for all but the smallest complexes in GE-PSN. Obviously, this high density in the inner-complexome is expected, where the biophysical and functional maps are detecting complexes, defined by independently curated experiments. Sys-NB-06 and GE-PSN are more enriched in the inter-complex pairs of Zone B. In Zones C and D, Sys-NB-06 and GE-PSN are depleted in regions involving the lower abundance proteins and enriched between the most abundant, with I3D-exp-20 and Lit-BM-20 showing a similar but less pronounced bias in their distributions, whereas ABBI-21, AlphaFold+RoseTTAFold, and GI-PSN are relatively uniformly distributed. This distribution of Sys-NB-06 and GE-PSN could be due to their dependence on endogenous expression. CS-PSN’s depletion in Zone C and enrichment in Zone D are dampened after correcting for the untested essential genes (Extended Data Fig. 5c-e). The uniform coverage of ABBI-21, AlphaFold+RoseTTAFold, and GI-PSN suggest that there are abundant biophysical and functional interactions between most of the proteome, regardless of expression levels.

**Fig. 5.**
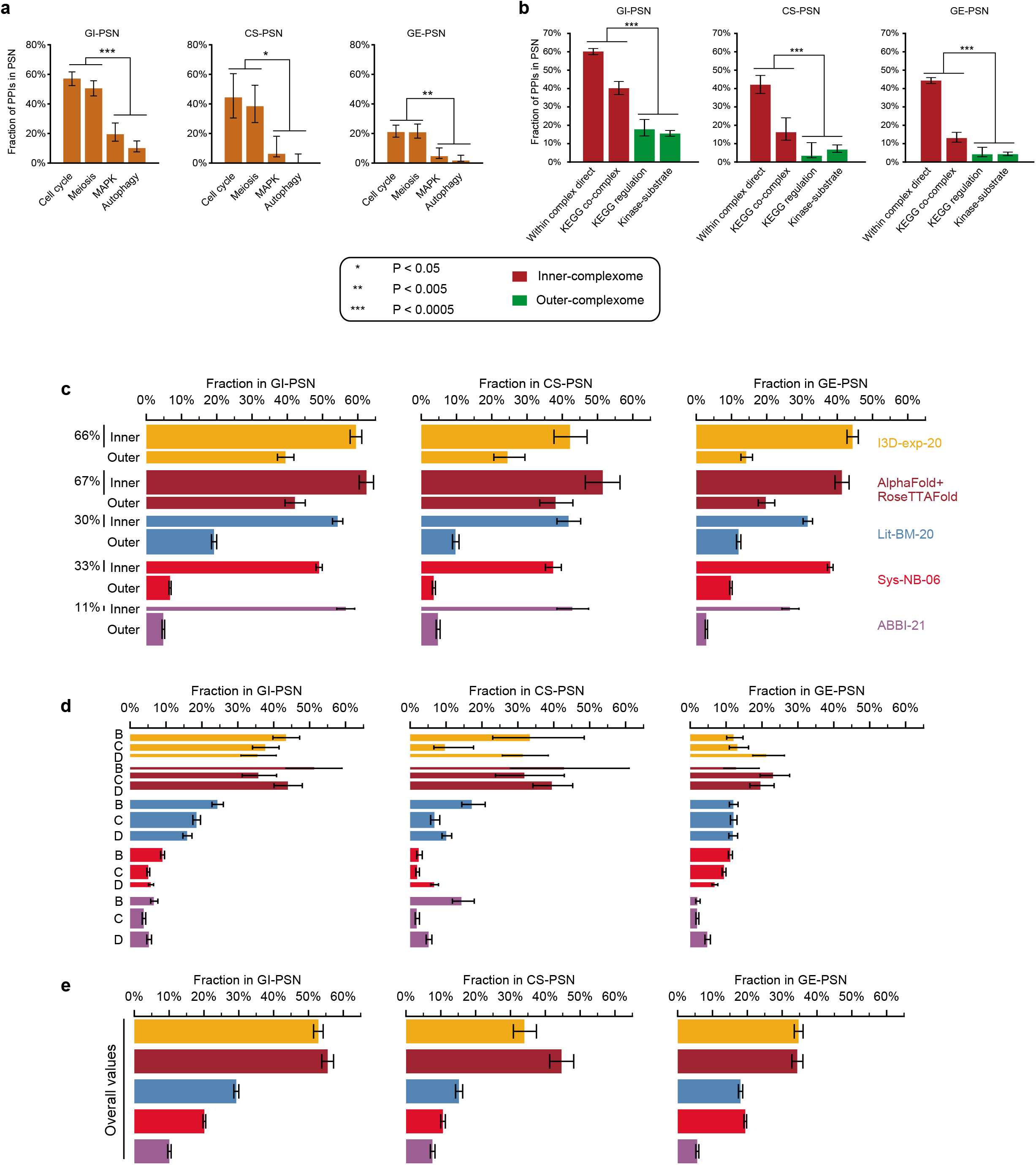
A large proportion of the interactome consists of interactions between functionally heterogeneous proteins in the outer-complexome. **a**, The fraction of PPIs from four different pathways connected in different functional networks. **b,** The fraction of PPIs from four gold-standard biophysical interaction datasets connected in functional networks. **c,** The fraction of pairs in the inner- and outer-complexome in the four biophysical maps connected in functional networks. The pie charts show the proportion of pairs in the inner- and outer-complexome within the four biophysical maps. **d,** The fraction of pairs from the outer-complexome, Zones B, C, and D, that are also connected in functional networks. **e** The fraction of pairs from four biophysical maps connected in functional networks. In all panels, error bars are 68.3% Bayesian credible intervals.

After extensively testing for bias across the different datasets, we found that systematic binary maps provide the most even coverage; however, one possible interpretation of this could be the presence of large numbers of randomly distributed false positives in the outer-complexome. To test whether ABBI-21 PPIs from either inner- or outer-complexome are of similar biophysical quality, we compared their recovery rates in MAPPIT and GPCA using Lit-BM-13 as a benchmark (Extended Data Fig. 5f-h). While ABBI-21 validates at a higher rate than Lit-BM-13 in the inner-complexome, PPI pairs from both ABBI-21 and Lit-BM-13 datasets show lower recovery rates in the outer-complexome than in the inner-complexome (*P* = 3 x 10^-12^, two-sided Fisher’s exact test). The fact that both our literature benchmark and ABBI-21 behave similarly in the outer-complexome demonstrates that ABBI-21 pairs in the outer-complexome are of good biophysical quality, suggesting that the difference in recovery rates between inner- and outer-complexome stems from differing biophysical factors, e.g. interaction affinity or post-translational modification dependency. Thus consistent with our previous observations that within-complex PPIs are detected more frequently in Y2H screens and that PPIs detected in more Y2H screens test positive in validation asssays at higher rates, independent of data quality^20^. The striking observation of inner-complexome PPIs being more readily detected by different PPI assays suggests that inner-complexome PPIs tend to be overrepresented in interactome maps relative to their proportion in the real interactome.

## Functional heterogeneity in the outer-complexome

To investigate how the difference between the inner- and outer-complexome relates to the functional relationships between physically interacting proteins, we computed the fraction of interacting protein pairs for which the corresponding gene pairs are also connected in the functional networks. We started by validating this approach using gold-standard PPIs from four well-characterised yeast pathways from the KEGG database: Cell cycle; Meiosis; MAPK signalling; and Autophagy^49^ (Fig. 5d). Interestingly, genes encoding interacting proteins from different pathways show large variation in the likelihood of being connected in the functional networks. Over half of the interacting pairs in Cell cycle and Meiosis pathways, essential for yeast growth and sexual reproduction, respectively, are connected in GI-PSN, whereas less than 20% of the interactions from the context-specific pathways – MAPK signalling and Autophagy – are detected (*P* = 0.002, two-sided Fisher’s exact test). GE-PSN and CS-PSN show a similar bias towards Cell cycle and Meiosis compared to MAPK and Autophagy (Fig. 5d). Next, we examined gold-standard reference PPIs from the inner- and outer-complexome^29, 49, 50^. All three PSNs captured significantly more interactions from the co-complex datasets than the outer-complexome PPIs in signalling pathways and kinase-substrate interactions (*P* ranges from 5 x 10^-9^ to 5 x 10^-45^, Fisher’s exact test) (Fig. 5e). A partial explanation for the detection of PPIs within constitutive rather than context-specific pathways is that GI-PSN was produced using yeast grown on rich media, in which environment the context-specific pathways will be mostly dormant, however, we also see differences in these pathways in CS-PSN and GE-PSN which each use data from a large number of different conditions. The most general explanation for these findings is that the PSNs are measuring functional similarity, at an aggregate level, of the two interacting proteins, with proteins that exist together in stable complexes being the most similar in their function, whereas proteins with transient interactions may often each have additional functions, independent from any specific binding partner, and so not be perfectly functionally similar at the aggregate level. This can be illustrated by the interaction between the importin-ɑ nuclear pore subunit Srp1 and the transcription factor Pho5. Although nuclear import is crucial for Pho5 function, the overall functions of both proteins are different and hence they are unconnected in the functional profile similarity networks.

After evaluating this approach using specialised sets of gold-standard PPIs, we next moved to the large-scale interactome maps, to evaluate the functional relationships between interacting proteins in the inner- and outer-complexome across the entire proteome. In all datasets, PPIs from the inner-complexome have a high probability of being connected in the functional networks and PPIs in the outer-complexome have lower probabilities (Fig. 5c, Extended Data Fig. 6). Interactions from all three zones of the outer-complexome, B, C, and D, within each biophysical map, are connected in the functional networks at similar rates, showing that the observed difference in the functional similarity of interacting proteins is a property of differences between the edges of the networks (PPIs) rather than between the nodes (proteins) (Fig. 5d). We observed consistent patterns when we investigated the specific examples of the ESCRT and OCA complexes (Supplementary Note 2, Extended Data Fig. 8). We observed similar patterns when directly using genetic interactions instead of the profile similarities. Pairs of genes encoding interacting proteins found in all biophysical maps have a higher likelihood to show negative than positive GIs^51^, and genes encoding protein pairs from the inner-complexome have an increased tendency to show negative GI than pairs in the outer-complexome (Extended Data Fig. 9a). The four biophysical datasets differ substantially in their proportion of inner-complexome pairs. More than one-quarter of pairs in Lit-BM-20 and Sys-NB-06, and more than half in AlphaFold+RoseTTAFold, are between subunits of the same complex (Zone A) compared to around one-tenth for ABBI-21 (Fig. 1d, 2i, 5b). This difference contributes to the lower aggregate fraction of ABBI-21 PPIs connected in functional PSNs, relative to Lit-BM-20 and Sys-NB-06, and the lower fraction for those three networks relative to I3D-exp-20 and AlphaFold+RoseTTAFold (Fig. 5e). Another factor affecting the overall rate of overlap with the different biophysical datasets is that GI-PSN is densest among essential genes^13^, and as a consequence, interactions between proteins encoded by essential genes show a higher likelihood to be connected in GI-PSN in all biophysical datasets. However, I3D-exp-20, AlphaFold+RoseTTAFold, Lit-BM-20, and Sys-NB-06 are also biased towards proteins encoded by essential genes, resulting in increased overlap with GI-PSN, whereas ABBI-21 covers the proteome and interactome more uniformly (Extended Data Fig. 9b). We cannot directly compare the rates of connection in the PSNs of the maps generated by systematic testing of large search spaces (Sys-NB-06 and ABBI-21) to those of the literature-curated datasets (Lit-BM-20 and I3D-exp-20), where most studies tend to focus on a particular pathway or process of interest. For example, a single study^52^ provides more than a quarter of the Lit-BM-20 pairs connected in GE-PSN by testing all pairs of 70 pre-ribosomal proteins, a test space with a density of 83% of pairs connected in GE-PSN compared to a density of 1% of pairs in the full proteome-by-proteome space connected in GE-PSN. Thus, the higher overlap of literature-curated PPIs with functional PSNs likely stems more from the choice of which protein pairs to test rather than the specific interactions detected, and the systematic maps should more accurately reflect the rate of connection in the PSNs for the real protein interactome. In summary, we find that the inner-complexome tends to consist of functionally similar interacting proteins. In contrast, the outer-complexome tends to consist of interactions between functionally heterogeneous proteins, presumably necessary for intracellular crosstalk.

One challenge to this interpretation is that I3D-exp-20 and AlphaFold+RoseTTAFold have high overlaps with functional networks in the outer-complexome relative to the other datasets. However, this could be because structures are biased towards permanent interactions, due to their ease of crystallisation relative to transient interactions, which often require additional techniques to crystallise^53^. To test this, we examined the interface area and predicted Δ*G* of the experimental and predicted PPI structures, finding that PPIs that are also connected in the functional networks tended to have larger interfaces and lower *ΔG* (Extended Data Fig. 7a, b, Supplementary Tables 18, 19). Computationally predicting structures should offer a way to overcome this bias in experimental data generation. However, by contrast we found that predicted structures had larger interfaces than the experimental structures (Extended Data Figure 6c), which was a result both of AlphaFold being more often able to generate a sufficiently confident predicted structure for larger interface PPIs (Extended Data Fig. 7d), and of a bias of AlphaFold predicting larger interfaces than seen in the experimental structures (Extended Data Fig. 7e). Together this suggests that, currently, AlphaFold+RoseTTAFold not only recapitulate the bias towards more stable PPIs in their training data but actually increase that bias in the PPIs for which they are able to generate confident predictions.

The structural networks exhibit truncated degree distributions, missing high degree, or hub, proteins (Extended Data Fig. 3i), possibly because of this bias against transient PPIs. Hubs are key features of PPI networks, which have a major impact on the network’s topology, and so we investigated the connectivity of PPI hubs in the functional PSNs. Hubs in biophysical maps can be classified as either ‘date’ or ‘party’ hubs depending on the degree to which interacting partners are also co-expressed^54^. Systematic binary maps have mostly date hubs, whereas literature-curated and systematic AP-MS maps have more party hubs^4^ (Extended Data Fig. 9c, d). Party hubs – which tend to be in complexes^55^ and use multiple interfaces to bind multiple partners simultaneously – overlap with functional networks twice as much as date hubs, which usually interact with partners one at a time transiently. The fraction of PPIs connected in the functional networks generally decreases as the interacting proteins’ degree increases across the different biophysical networks (Extended Data Fig. 9e). One notable exception is Sys-NB-06 in GE-PSN due to the tightly correlated expression of large protein complexes’ subunits, and the inclusion of pairs corresponding to indirect associations in those complexes. AlphaFold+RoseTTAFold displays very different trends, probably because of its truncated degree distribution. The observation that high-degree proteins are less functionally similar to their binding partners is consistent with the observation that they are more pleiotropic^4^.

## Discussion

Four decades ago, McConkey proposed the term “*quinary structure*” as a “fifth level of organization”, referring to macromolecular interactions that, although potentially highly functionally relevant, might be “transient *in vivo”*^56–58^. He predicted such interactions “will not be evident from the composition of purified proteins”, since, while quaternary structures tend to be more resistant to the “cataclysmic violence of the most gentle homogenization procedure”, quinary interactions, “although stable *in vivo*, might be largely destroyed by cell fractionation”. The observations presented in this paper suggest fundamental differences in organisation between the inner-complexome, containing mostly quaternary structures that are highly detectable by affinity purification approaches, and the outer-complexome, which has a greater tendency towards quinary structures, detectable by binary assays in living cells. From this perspective, it is easy to see why there should be a substantial discrepancy among alternative views of inner- versus outer-complexome organisation revealed by approaches as different as co-complex association detection and binary interaction assays. Indeed, we observe that the different methods used to obtain PPIs have different propensities to include transient interactions as well as different proportions of direct interactions and indirect co-complex associations, and that these differences can have a dramatic influence on the properties of the resulting network and hence our understanding of cellular organisation.

Although undoubtedly oversimplistic, our results evoke a view in which the inner-complexome constitutes the “manufacturing machinery” operating in a relatively constant, robust, and persistent manner, and the outer-complexome comprises the “regulatory processes’’ exhibiting greater flexibility, plasticity, environmental responsiveness, and evolvability. A relatively small proportion of the interactome is composed of inner-complexome interactions, while the vast majority of the interactome consists of interactions in the outer-complexome, consistent with emerging evidence suggesting that the majority of the interactome may be transient and context-specific^25–27^, between complementary rather than similar proteins^59^, and with extensive pathway cross-talk and pleiotropy^60^. However, mapping and functional characterization of these outer-complexome interactions remain challenging. Continued development of assays that can efficiently detect transient, context-specific PPIs, combined with new large-scale approaches to characterize their functions will be an important next step toward understanding the global organisation of cellular processes.

## Acknowledgements

We thank S. de Rouck for help with the MAPPIT experiments. We thank Gary Bader and acknowledge past and current members of the Center for Cancer Systems Biology (CCSB) for helpful discussions and experimental help. This work was supported primarily by National Institutes of Health (NIH) grant R01HG006061 (M.V, D.E.H, M.A.C, M.E.C, P.F.-B.) with additional support from R01GM130885 (M.V.), R01GM133185 (M.V., M.A.C, F.P.R.), and Institute Sponsored Research funds from the Dana-Farber Cancer Institute Strategic Initiative (M.V). A.D. was supported by a Leon Fredericq grant and a FRS-FNRS-Télévie Fellowship #7651317F (J-C.T). M.V. is a Chercheur Qualifié Honoraire from the Fonds de la Recherche Scientifique (FRS-FNRS, Wallonia-Brussels Federation, Belgium). J-C.T is Maitre de Recherche of the FRS-FNRS. F.P.R was supported by a CIHR Foundation grant. D.-K.K. was supported by a Banting Postdoctoral Fellowship through the Natural Sciences and Engineering Research Council (NSERC) of Canada and by the Basic Science Research Program through the National Research Foundation (NRF) of Korea funded by the Ministry of Education (2017R1A6A3A03004385). M.W.M received support from the Google Summer of Code 2015 through the National Resources For Network Biology (NRNB). C.P. was supported by a Ramon y Cajal fellowship (RYC-2017-22959). A.Y. was supported by a Deborah F. Allinger Fellowship awarded to CCSB.

## Author contributions

Computational analyses were performed by L.L., Y.W., with help from B.C., D.D.R., T.R., and K.L. Interactome mapping experiments were performed by A.D., T.C., with help from K.S., S.S., N.J., Q.Z., Z.Y., and K.S. Sequencing to identify interacting proteins was carried out by A.G.C., M.G., N.K., J.J.K., and J.C.M. Y2H vectors were designed and generated by Q.Z. with help from N.J. The preparation of Y2H, GPCA, and MAPPIT destination clones by *en masse* gateway cloning and yeast transformations were performed by Q.Z., N.J., A.D., and T.C. Experimental results were processed by Y.W., T.H., and K.L. GPCA validation experiments were done by A.D., and T.C., with help from Y.J. MAPPIT validation experiments were done by I.L., supervised by J.T. Functional enrichment analyses were done by D.-K.K., L.L., and Y.W. Extraction of the literature datasets was performed by L.L., and T.H. YeRI web portal was built by M.W.M., supervised by J.R., and M.H. Structural analyses were done by C.P., L.L., and Y.W., supervised by P.A. Topological analyses were done by L.L. Sequencing analyses were done by T.H., W.B., Y.S., and Y.W. Network-based functional prediction was performed by I.K. The overall research effort was designed and conceptualised by M.V., F.P.R., M.A.C., D.E.H., P.F.-B., Y.W., A.D., L.L., and A.Y. Interactome mapping was supervised by B.C., M.V., M.A.C., D.E.H., and T.H. Draft manuscript was written by L.L., A.Y., Y.W., A.D., M.V. Manuscript was reviewed and edited, with contributions from other co-authors, by L.L., A.Y., Y.W., M.V., D.E.H, T.H., J.-C.T., F.P.R., and M.A.C. The overall research effort was supervised and/or advised by M.V., F.P.R., M.A.C., D.E.H. The project was conceived by M.V. Major funding acquisition was by M.V., D.E.H., M.A.C., F.P.R. P.F.-B., M.E.C., and J.-C.T.

## Materials and Correspondence

should be addressed to D.E.H., M.A.C., J.-C.T, or M.V.

## Data availability

YeRI, ABBI-21, and Lit-BM-20 maps are available at our OpenPIP^61^ website: http://yeast.interactome-atlas.org/.

## Code availability

Analysis code and data are available at https://github.com/ccsb-dfci/YeRI_paper.

---

**Extended Data Fig. 1.**
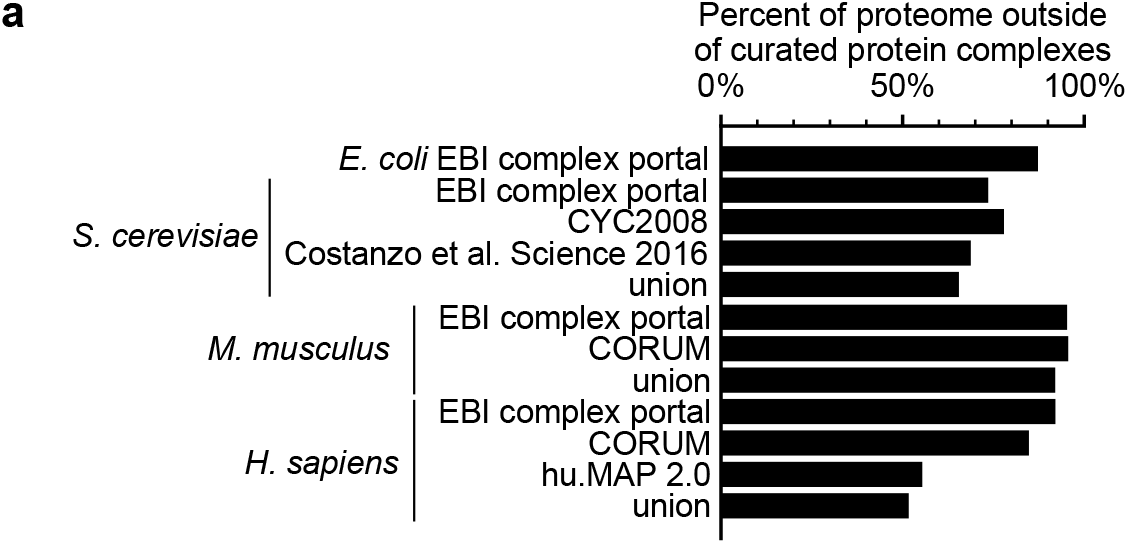
Most proteins are not members of protein complexes. **a**, The fraction of the proteome of different organisms that are not listed as subunits of protein complexes of at least 3 or more different protein subunits, using different datasets of protein complexes. The union of the protein complex datasets for each organism is also shown, when there is more than one dataset.

**Extended Data Fig. 2.**
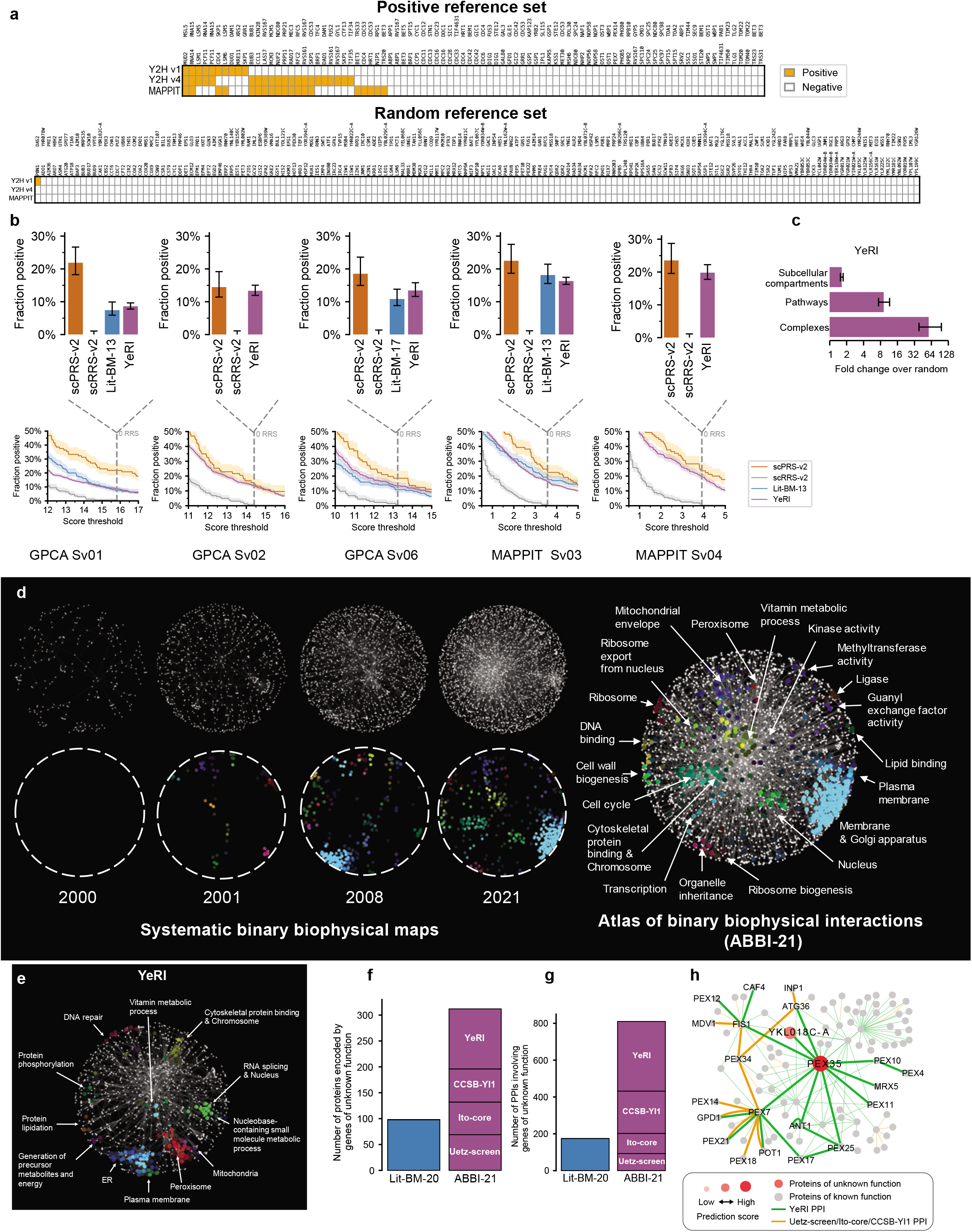
Validation of YeRI. **a**, Benchmarking Y2H v4 against two other binary PPI assays, using positive and random reference sets (scPRS-v2 and scRRS-v2). Coloured bars indicate positives. **b**, The results of different batches of experimental validation of YeRI in GPCA and MAPPIT. In total, all pairs from YeRI were tested in both assays. Error bars and shaded bands are 68.3% Bayesian credible intervals. **c**, Enrichment of YeRI PPIs between proteins in the same cellular compartments, pathways, and protein complexes. Error bars are 68.3% interval of degree-preserved random networks. **d**, Composite networks, generated by the addition of each systematic PPI map (top row, left). Network regions enriched for GO terms (bottom row, left). Merge of network and enriched regions for the most recent composite network (right). **e**, Network-based spatial enrichment analysis (SAFE) for YeRI. Clusters of genes enriched for GO terms in YeRI are highlighted. **f**, The number of proteins encoded by genes of unknown function with at least one interaction in ABBI-21 or Lit-BM. **g**, Number of PPIs involving proteins encoded by genes of unknown function in Lit-BM-20 or ABBI-21. **h**, PPI network of PEX35 and its first- and second-degree interactors. Named proteins are those annotated or predicted to have the GO terms related to ‘peroxisome importer complex’.

**Extended Data Fig. 3.**
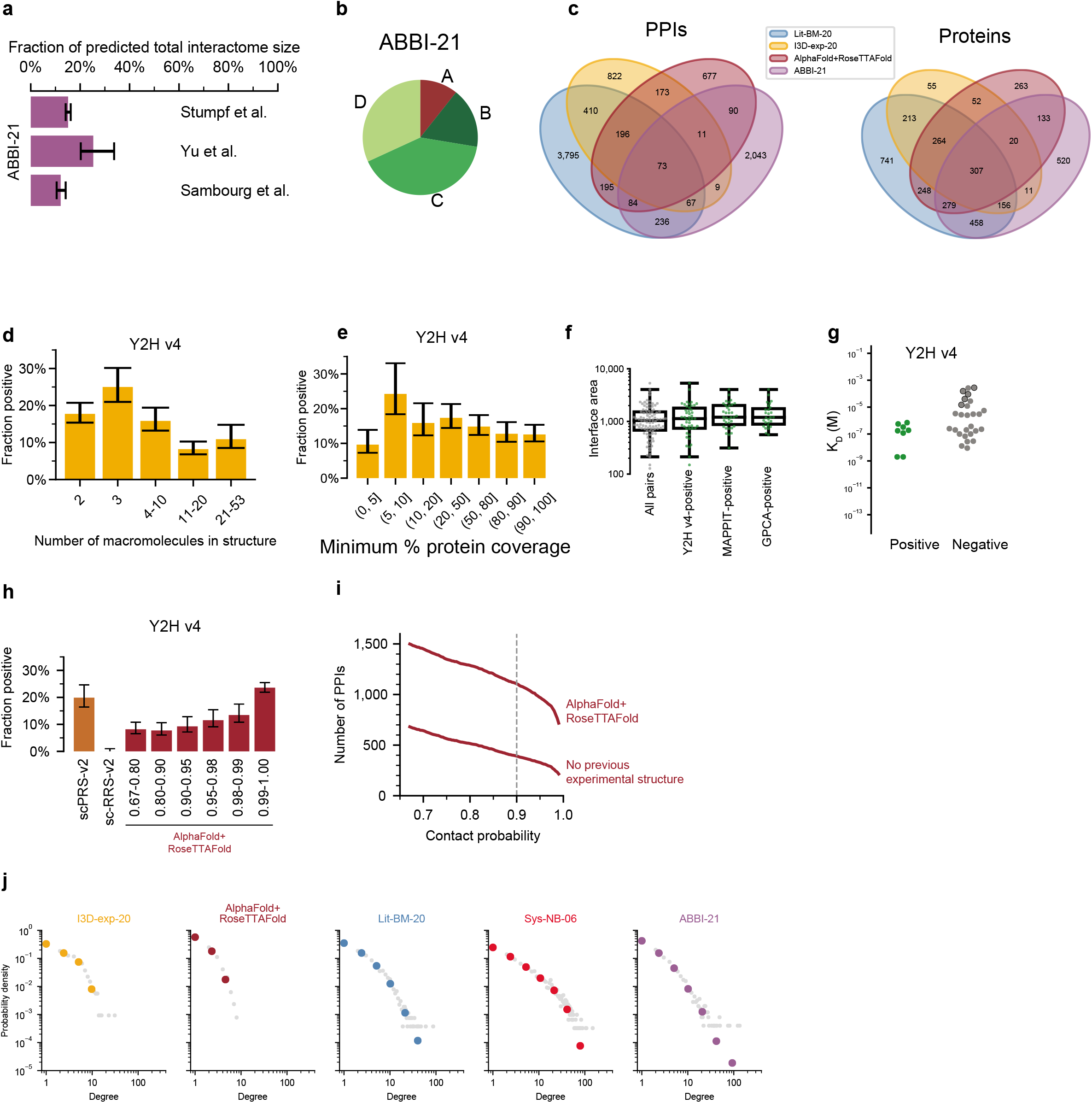
Assessment of high-quality binary PPI datasets. **a**, Coverage of the yeast interactome by ABBI-21, based on three reported estimates of the total interactome size. Error bars correspond to the 95% confidence intervals of each estimate reported in the original publications. **b**, The proportion of ABBI-21 in each Zone. **c**, Venn diagram of PPIs, and proteins with at least one PPI, in four high-quality binary datasets. **d**, Fraction of PPIs identified by Y2H v4 in complex structures of different sizes. Error bars are 68.3% Bayesian credible intervals. **e**, Y2H v4 recovery of I3D-exp-17 pairs as a function of the fraction of the protein that is contained in the experimental structure. The lower of the two values of coverage of the different proteins in each interaction is used. Error bars are 68.3% Bayesian credible intervals. **f**, Distribution of interface area for I3D-exp-17 PPIs, that tested positive in Y2H v4, MAPPIT, and GPCA, restricting to pairs that were successfully tested in all three assays. Box plots show median, interquartile range (IQR), and 1.5×IQR. **g**, Dissociation constants of pairs positive or negative in Y2H v4. Points outlined in black are PPIs (non-covalent interactions) of different ubiquitin-binding proteins with ubiquitin. **h**, Results of testing AlphaFold+RoseTTAFold in Y2H v4, split into bins of contact probability. Error bars are 68.3% Bayesian credible intervals. **i**, Number of PPIs in AlphaFold+RoseTTAFold dataset as a function of contact probability. The lower line shows the number of PPIs without a previous experimental structure of the complex, made with either the exact *S. cerevisiae* proteins or homologous proteins. **j**, degree distribution of each network.

**Extended Data Fig. 4.**
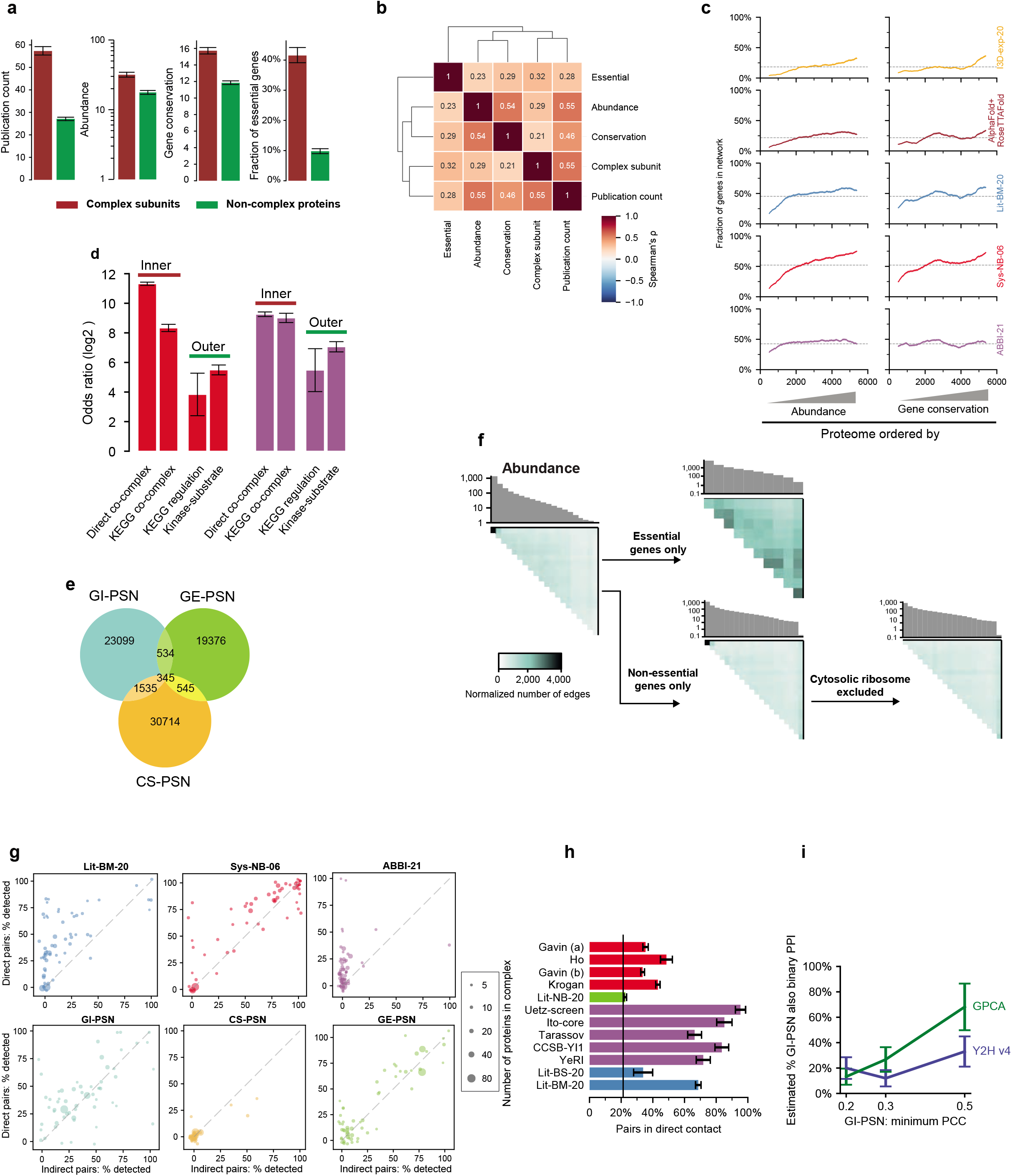
Validation experiment results in different Zones. **a**, Mean of publication count, abundance, gene conservation, and the fraction of essential genes for proteins either in or outside complexes. Error bars are 95% confidence interval. **b**, Correlation matrix with clustering of different protein-level properties based on Spearman’s rank correlation coefficient. **c**, The fraction of genes in five biophysical networks across sliding windows of 1,000 proteins, ordered by abundance and conservation. Grey dotted lines show the overall fraction. **d**, Enrichments of Sys-NB-06 and ABBI-21 to contain different gold-standard sets of PPIs from the inner- and outer-complexome. Direct PPIs within protein complexes with three or more subunits; co-complex pairs from KEGG pathways; PPIs regulating activation or inhibition from KEGG; and high-quality kinase-substrate pairs from the KID database. Error bars are standard error on the log odds ratio. **e**, Overlap of the three functional profile similarity networks (PSNs), genetic interaction (GI-PSN), gene expression at mRNA level across different conditions (GE-PSN), and growth across different conditions PSN (CS-PSN). Restricted to genes tested in all three PSNs with PCC value in the top 1% of tested pairs in each PSN. **f**, Investigation of densely connected area between the most intensely studied genes in GI-PSN. Heatmaps of the number of connected gene pairs in GI-PSN, ordered by protein abundance, and further segmented based on essentiality and involvement in the ribosome. **g**, Recovery of protein pairs in direct contact and pairs not in direct contact within the corresponding 3D structures, by different biophysical and functional networks. Each point is a separate protein complex with at least 5 distinct protein subunits. **h**, The fraction of directly contacting interactions, taking all reported pairs where both proteins are in a protein complex structure with at least three subunits, for different datasets. Error bars are standard error of proportion. **i**, Estimation of the fraction GI-PSN pairs that are also a binary PPI, based on results of testing samples from GI-PSNs at different PCC cutoffs using Y2H v4 and GPCA and comparing to the results of the PRS. Error bars are 1σ confidence intervals.

**Extended Data Fig. 5.**
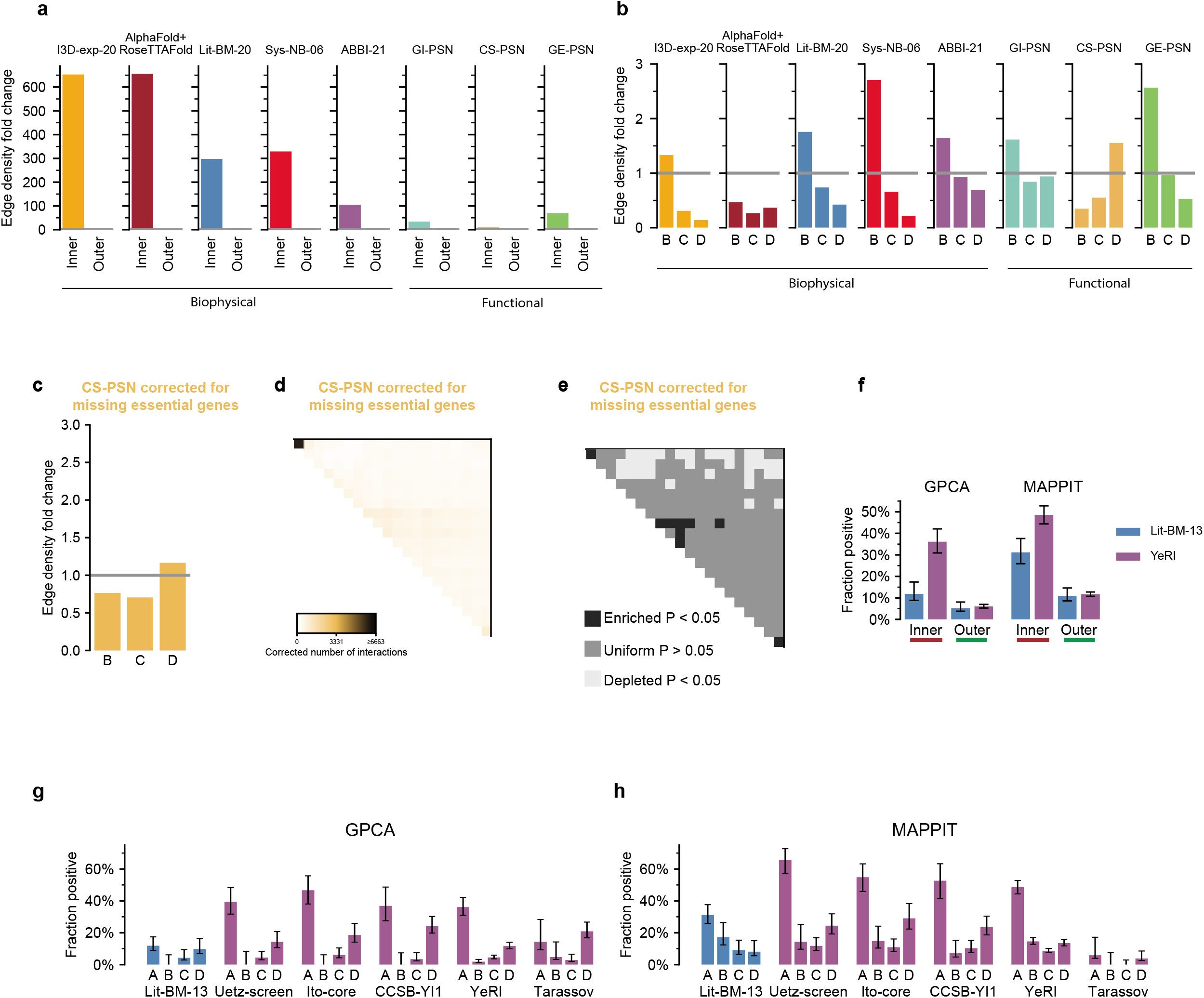
A large proportion of the interactome mainly consists of interactions between functionally heterogeneous proteins in the outer-complexome. **a**, Edge density fold change in the inner- and outer-complexome in the biophysical and functional maps. **b**, Edge density fold change in the Zones B, C and D of the outer-complexome in the biophysical and functional maps. **c**-**e**, Distribution of CS-PSN after restricting to genes tested in the experiment: **c**, edge density fold change in the outer-complexome (Zones B, C and D), **d,** heatmap of the number of connected gene pairs with the proteome first ordered by size of protein complexes in which the proteins are involved then ordered by protein abundance, and, **e**, statistically significant enrichment and depletion of pairs in **d**. **f**, Fraction of pairs in the inner- and outer-complexome from Lit-BM-13 and ABBI-21 that tested positive using GPCA and MAPPIT. **g**, **h**, The fraction of pairs testing positive in GPCA (**g**) and MAPPIT (**h**) in Zones A, B, C, and D from Lit-BM and systematic binary maps. All error bars are 68.3% Bayesian credible intervals.

**Extended Data Fig. 6.**
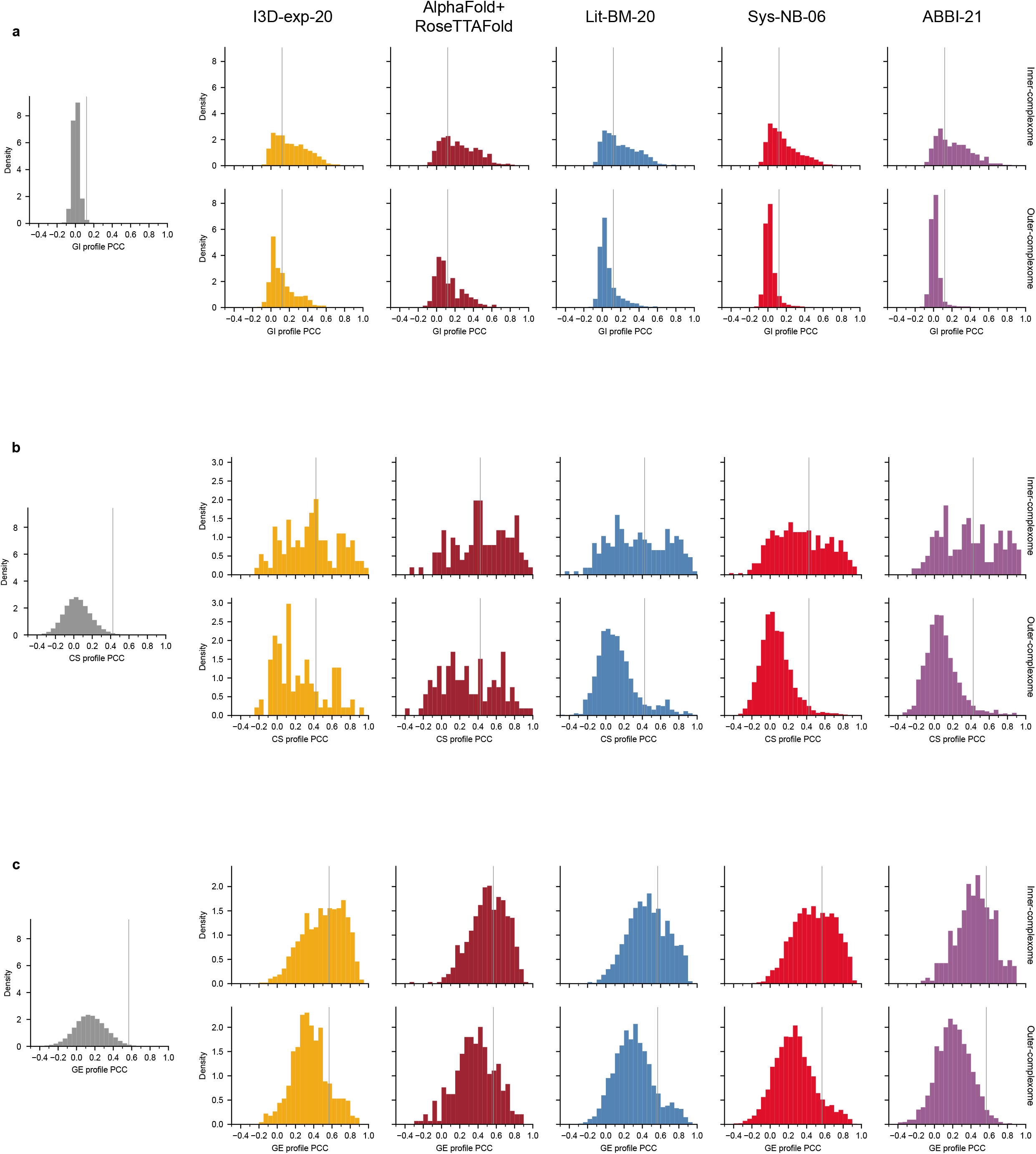
Distributions of functional profile similarity values of physically interacting proteins produce consistent results with the overlap with PSNs analysis. **a**, **b**, **c**, Left panel: PCC value for each of the functional profiles across all tested gene pairs. Right panels: distribution of functional profile PCC for interacting proteins, in each PPI network, split into inner- and outer-complexome. Grey vertical lines correspond to the cutoff used to make each PSN. GI-PSN (**a**), CS-PSN (**b**), GE-PSN (**c**).

**Extended Data Fig. 7.**
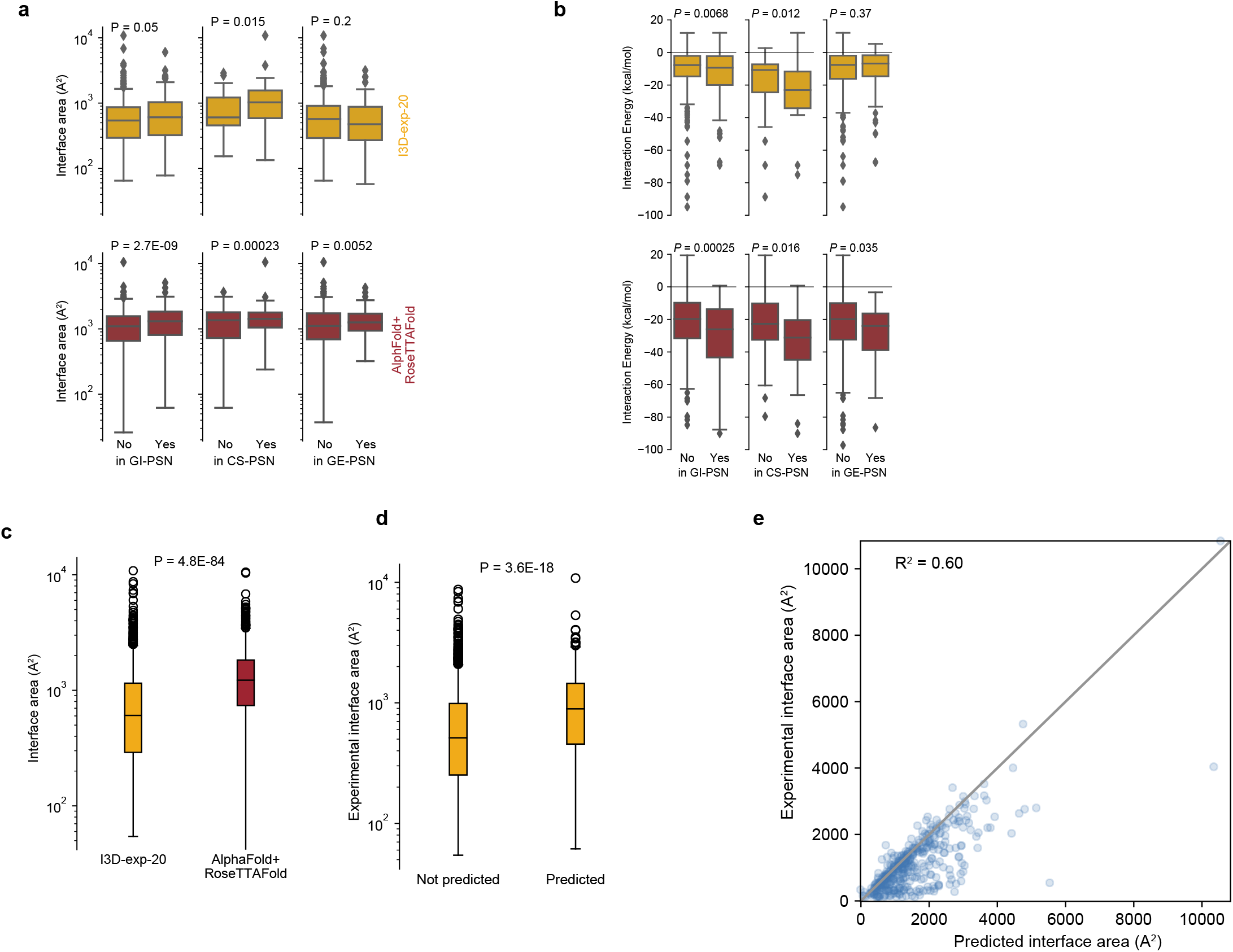
Stronger PPIs more often overlap with functional similarity networks. **a**, **b**, Distribution of PPI interface sizes (**a**), and predicted Δ*G* of interaction (**b**), from experimentally solved and computationally predicted structures, comparing PPIs that are connected in functional PSNs to those that are not. **c**, Distribution of PPI interface sizes between the experimental and predicted structure datasets. **d**, Distribution of PPI interface sizes, using the experimental structures, split by whether the PPI appears in the predicted structures dataset. All box plots show median, interquartile range (IQR), and 1.5×IQR (with outliers). **e**, Comparison of the PPI interface size of the computationally predicted structure to the experimental structure, for PPIs with both predicted and experimental structures. *P*-values in all panels calculated using two-tailed Mann-Whitney U test.

**Extended Data Fig. 8.**
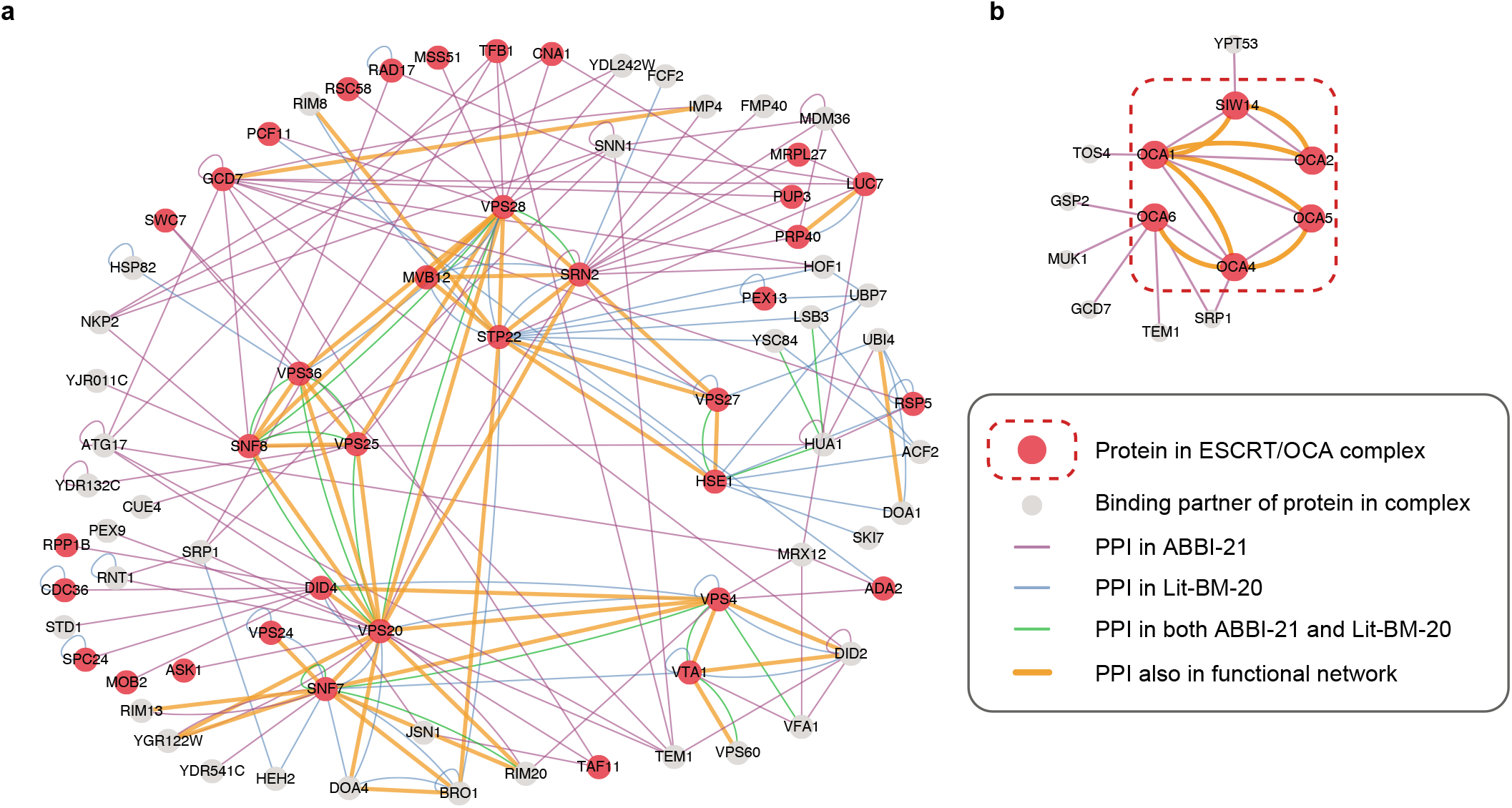
ESCRT / OCA complexes. **a**, Integrated network showing interactions among subunits of ESCRT complexes and their interacting partners. **b**, Integrated network showing interactions among subunits of the OCA complex and their interacting partners. Physically interacting proteins also connected in functional networks are connected with thick orange lines.

**Extended Data Fig. 9.**
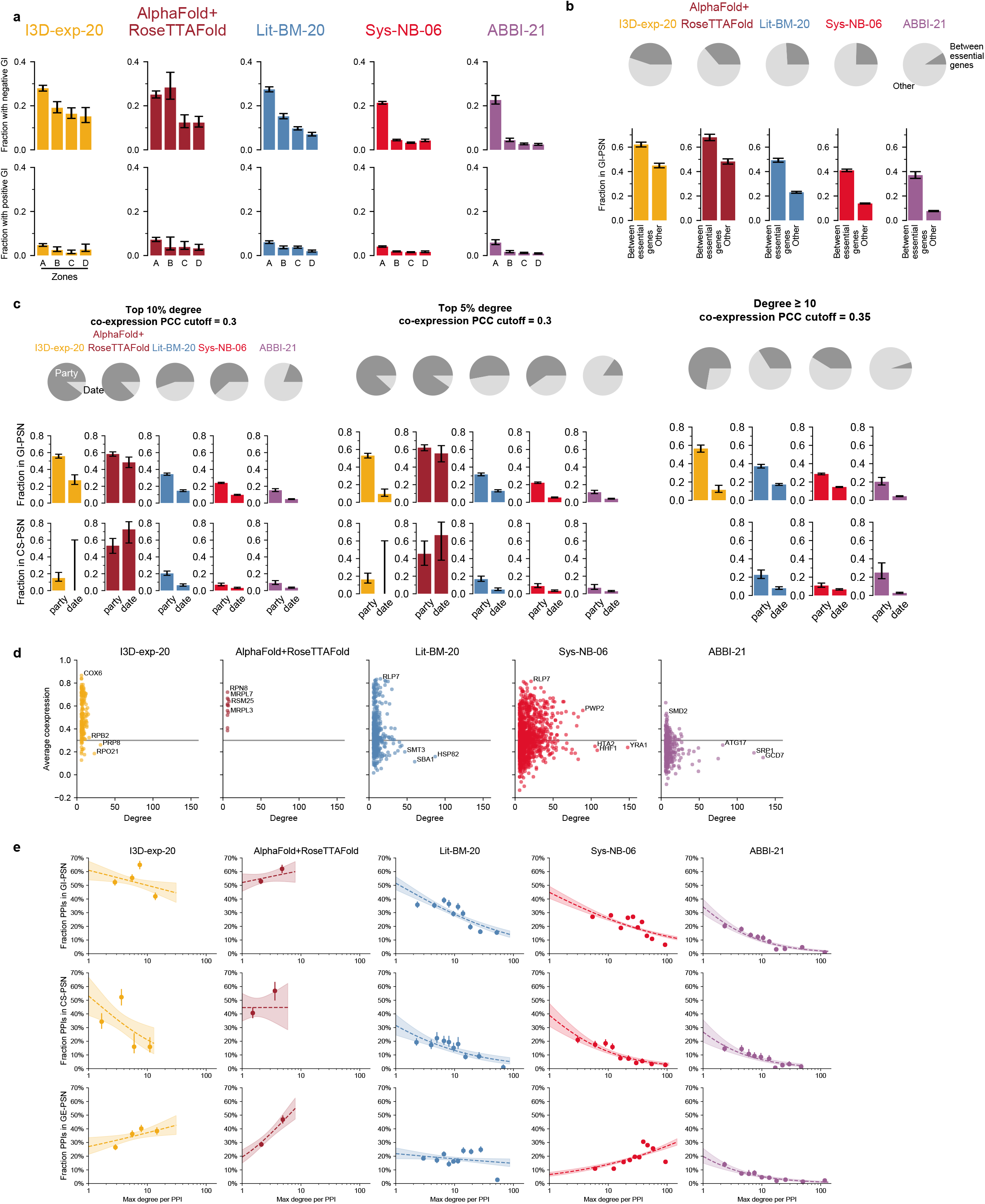
Further exploration of factors affecting PPI connection in functional networks. **a**, Composition of edges from three biophysical maps in inner-complexome (Zone A) and outer-complexome (Zones B, C, and D) and the fraction of pairs from biophysical maps also connected in positive or negative genetic interaction networks (top 1%) within the four zones. Error bars are 68.3% Bayesian credible intervals. **b,** Composition of edges from three biophysical maps between essential genes and involving non-essential genes, and the fraction of pairs from biophysical maps of each category connected in functional networks. Error bars are 68.3% Bayesian credible intervals. **c**, Composition of edges from three biophysical maps involving date and party hubs using different degree and PCC thresholds, and the fraction of pairs from biophysical maps of each category connected in functional networks. Only panels for which there are at least one date and one party hub are shown. Error bars are 68.3% Bayesian credible intervals. **d**, Average co-expression PCC of proteins in biophysical maps with different degrees. A cutoff between party and date hubs of 0.3 is shown by the grey horizontal line. **e**, The fraction of edges that are also connected in functional networks, for three biophysical maps, binned by the higher degree of the two proteins for each pair. Logistic regression and 95% confidence interval are shown.

## Supplementary Information

### Supplementary Note 1 – Predicting functions for genes with YeRI

Despite being one of the most well-studied organisms, the function of almost one-sixth of yeast genes remains unknown ^63^. We investigated the number of PPIs involving the products of these uncharacterised genes in both literature and systematic maps. Systematic maps identify substantially more PPIs connecting proteins encoded by genes of unknown function than literature-derived maps (Extended Data Fig. 2g). Altogether, 33% of such proteins have at least one interaction in ABBI-21, while 19% have interactions identified only in YeRI (Extended Data Fig. 2f). Given the lack of progress in characterizing the functions of these genes, systematically mapped PPIs provide information to infer their cellular roles. We predicted functions for genes of unknown function with a guilt-by-association approach using GO term annotations of their interaction partners^20^ (Supplementary Table 13) in ABBI-21. The lag between the release of publications describing gene functions and the curation of that information into GO terms sometimes results in a small number of genes which appear in the GO-term based list of genes as “genes of unknown”, while in fact, they have been assigned functions already. These cases present an opportunity to test the accuracy of our predictions. For example, a gene of unknown function *YGR168C* is also known as *PEX35* owing to its recently demonstrated role as a regulator of peroxisomal abundance ^64^. Using ABBI-21, we predicted *YGR168C* to be involved in peroxisomal protein import machinery, showcasing the ability of ABBI-21 to accurately predict gene function. Pex35 has 23 PPIs in ABBI-21, all from YeRI, out of which eight are proteins involved in peroxisomal biology (Extended Data Fig. 2h). Pex35 also interacts with another protein encoded by a gene of unknown function, *YKL018C-A*/*MCO12*, which we predict to be involved in peroxisomal abundance as well. Another example of the efficacy of using a guilt by association approach with our systematic PPI network to predict gene function is *YJR015W*, the product of which was recently demonstrated to be localised to the ER ^65^. Indeed, we predicted *YJR015W* as a putative facilitator of endoplasmic reticulum (ER) transport activity, based on its protein interactions with ER secretory pathway components such as Sec11, Spc1, and Sar1.

### Supplementary Note 2 – ESCRT / OCA complexes

The endosomal sorting complex required for transport (ESCRT) pathway plays a key role in the biogenesis of multivesicular bodies and turnover of membrane proteins^66^. The main players in the ESCRT pathway are the five ESCRT complexes, supporting auxiliary proteins, and the cargo to be sorted. By integrating biophysical and genetic networks, we observe that the five ESCRT complexes’ core constituents interact biophysically in both ABBI-21 and Lit-BM-20 and are highly interconnected in functional networks (Extended Data Fig. 8a). In contrast, outer-complexome ABBI-21 PPIs between subunits of ESCRT complex and non-complex proteins, important for endosomal sorting, are not connected in the functional networks. For example, ABBI-21 contains PPIs between Vfa1, important for vacuolar sorting, and Vps4 and Vta1, subunits of the ESCRT-4 complex. However, despite known functional roles, Vfa1 is not connected with Vps4 and Vta1 in any of the PSNs.

Systematic binary maps can help us understand how proteins within and outside complexes function together to mediate various biological processes. One such example is Snn1, a subunit of the biogenesis of lysosome-related organelles complex 1, BLOC-1, important for endosomal maturation^67, 68^. In ABBI-21, Snn1 interacts with proteins of the ESCRT complex like Vps28 and other non-complex endosomal proteins like Nkp2 (Extended Data Fig. 8a). ABBI-21 interactors of Snn1 are significantly enriched in proteins located in endosomes (13%, vs 2% overall for proteins in ABBI-21, *P* = 0.0007, two-sided Fisher’s exact test). Five out of six BLOC-1 complex proteins have PPIs primarily in ABBI-21, and none of the interacting protein pairs are connected in any of the functional networks.

The uniform coverage of inner- and outer-complexome by ABBI-21 can also shed light upon potential mechanisms by which previously under-studied complexes act. For example, the oxidant-induced cell-cycle arrest (OCA) complex mediates G1 arrest under stress conditions through a yet unknown mechanism^69^. This complex’s six components are connected biophysically and in the functional networks, exhibiting similar genetic interaction and condition sensitivity profiles (Extended Data Fig. 8b). Although inner-complexome interactions with OCA may well be critical for its function, they do not explain the complex’s stress-specificity. Outer-complexome interactions of OCA proteins do not overlap with the genetic networks but might be instrumental in understanding the mechanism through which the complex mediates its function. Of particular interest is the interaction between Oca1 and Tos4, newly reported in YeRI (Extended Data Fig. 8b). Tos4 is a transcription factor that binds to the promoters of genes involved in the G1/S transition^70^, offering a hypothesis for the mechanism by which OCA mediates G1 arrest.

### Methods

#### Strains and cell lines

##### Yeast strains

Yeast haploid strains *MAT*α Y8930 and *MAT***a** Y8800, derived from PJ69-4^71^, were used previously^4, 72^. Both strains harbor the following genotype: *leu2-3,112 trp1-901 his3Δ200 ura3-52 gal4Δ gal80Δ GAL2::ADE2 GAL1::HIS3@LYS2 GAL7::lacZ@MET2 cyh2^R^*. Yeast cells, parental strains or transformants, were cultured either in YEPD or synthetic drop out media, supplemented as needed and incubated at 30°C.

##### Bacterial strains

Chemically competent DH5α or DB3.1 *E. coli* cells were used for all bacterial transformations in this study. Transformed cells were cultured in Luria Broth or Terrific Broth, supplemented with antibiotics (50 µg/ml of ampicillin, spectinomycin or kanamycin) as needed and incubated at 37°C.

##### Human cell lines

Human embryonic kidney HEK293T cells were cultured in Dulbecco’s Modified Eagle Medium (DMEM) supplemented with 10% fetal bovine serum, 2mmol/L L-glutamine, 100 I.U./mL penicillin, and 100 μg/mL streptomycin. Cells were incubated at 37°C with 5% CO2 and 95% humidity.

#### Yeast open reading frames

The list of yeast ORFs was downloaded from the *Saccharomyces* Genome Database (SGD) (https://www.yeastgenome.org/) on January 14^th^, 2017. Four ORFs (YCR097W/HMRA1, YCR096C/HMRA2, YCL066W/HMLALPHA1, YCL067C/HMLALPHA2) annotated in SGD as “silenced gene” were removed. Only SGD-annotated “Verified” and “Uncharacterized” ORFs were included whereas ORFs annotated as “Dubious” were excluded, leaving a total of 5,883 ORFs with 5,155 and 728 ORFs classified as Verified and Uncharacterized, respectively. All datasets analyzed have been restricted to these 5,883 ORFs and previous ORF names that appear as aliases for one of these ORFs have been mapped to their corresponding new name.

#### Complexome – list of protein complexes

Most analyses use yeast complexes taken from Data File S12 of Costanzo *et al.* 2016^13^ and filtered to contain three or more different protein subunits, resulting in 339 complexes containing 1,897 different proteins. For the cross-species analysis in Figure 1a, the data came from the EBI Complex Portal^73^ dated Feb 3^rd^ 2022, and also filtered for those containing three or more different protein subunits. Additional datasets were used in Extended Data Figure 1a, from CYC2008^45^, CORUM^74^ v3 dated 3^rd^ September 2018, and Hu.MAP 2.0^75^ dated 9^th^ August 2020, all filtered to contain three or more different protein subunits.

#### Assigning protein pairwise combinations to individual zones

The search space of all possible pairwise combinations of proteins can be classified into four different “zones” based on their relationship to the complexome (Figure 1A). We define Zone A, which we refer to as the inner-complexome, as all pairwise combinations of proteins within protein complexes. Such pairs would include for example Rpt4 and Rpt5, two interacting subunits of the proteasome^76^, and Rps1A and Rps14A of the ribosome^77^. Zone B corresponds to pairs of proteins where each protein belongs to a different complex. For example, the RNA polymerase II (RNA Pol-II) Rpb2 subunit is capable of interacting with the Tfg2 subunit of the transcription factor II complex TFIIH^78^. Zone C represents all pairwise combinations where one protein is in a complex and the other is not. For example, Rpl10, a component of the large ribosomal subunit, interacts with Sqt1, a chaperone important for Rpl10 assembly into the ribosome^79^. Another example would be Rbp2 which interacts with Rad26, a nucleotide excision repair protein recruited to DNA lesions by RNA Pol II^80^. Finally, Zone D corresponds to protein pairs where neither protein belongs to a complex. Examples of Zone D interactions include most PPIs within signal transduction pathways, individual chaperones and their clients or kinase-substrate pairs involved in cellular processes such as autophagy.

While populated by relatively abundant proteins and large molecular size machines, the inner-complexome covers only a tiny proportion of the full yeast interactome “search space”, *i.e.* all ∼18,000,000 pairwise combinations between all ∼6,000 proteins. For example, the yeast ribosome, which accounts for nearly 20% of the proteomic mass^62^, is encoded by only 2% of all genes and all combinations between ribosomal proteins correspond to ∼0.04% of the whole search space. Together the 339 complexes in our complexome map represent 17,607 pairwise combinations between their respective subunits, which corresponds to only ∼0.1% of the proteome-by-proteome space. This leaves us with ∼99.9% of the whole search space for the outer-complexome, with its three zones, B, C, and D, corresponding to 10%, 44%, and 46% of the proteome-by-proteome space, respectively.

#### Assembly and description of biophysical and genetic datasets

##### Y2H-union: Uetz-screen, Ito-core and CCSB-YI1

As described previously^4^, Uetz-screen is a subset of PPIs from Uetz et al^2, 3^ that was obtained from a proteome-scale systematic Y2H screen, excluding a smaller-scale, relatively biased, targeted experiment with a smaller number of well-studied bait proteins. Ito-core is a subset of PPIs found three times or more in Ito et al^3^, excluding unreliable pairs of proteins found only once or twice. CCSB-Y1 is a proteome-scale dataset of Y2H PPIs validated using the two orthogonal assays MAPPIT and yPCA^4^. After restricting to PPIs involving the 5,883 ORFs (described above) the dataset sizes are as follows: Uetz-screen: 645 PPIs; Ito-core: 816 PPIs; CCSB-YI: 1,772 PPIs. The union of these maps (Y2H-union) contains 1.933 nodes and 2,833 PPIs.

##### Literature-curated biophysical datasets (Lit-NB, Lit-BS, Lit-BM)

Literature-curated pairs were obtained from the databases MINT^9^, IntAct^8^, DIP^10^, and BioGRID^11^. The data files used were the 2020-07-14 release from IntAct (containing data from IntAct, MINT and DIP) and BioGRID release 3.5.187 (from 2020-06-25). We excluded evidence corresponding to the eight systematic, proteome-scale co-complex association datasets^2–7, 38, 46^. Data was filtered to ensure valid IDs for UniProt accession numbers, Pubmed IDs and PSI-MI terms. Each piece of evidence for a protein pair had to consist of a Pubmed ID and an interaction detection method code in the PSI-MI controlled vocabulary (http://www.psidev.info/groups/molecular-interactions). Duplicated evidence can arise in cases where different source databases curate the same paper. We merged duplicated entries for each pair, as detected by multiple pieces of evidence with the same Pubmed ID and experimental interaction detection codes which are either identical or have an ancestor-descendent relationship in the PSI-MI ontology. In the latter case, the more specific descendent term was assigned to the merged evidence. In order to select the subset of protein pairs corresponding to binary interactions (as opposed to co-complex associations), we developed a manual classification of the PSI-MI interaction detection method terms^39^. Our classification has since been updated to cover new experimental methods which have been added to the controlled vocabulary in the intervening time. The methods are classified into three categories; ‘invalid’, ‘binary’ and ‘non-binary’. Where ‘invalid’ corresponds to PSI-MI terms that are not considered valid experimental protein-protein interaction detection methods, ‘binary’ corresponds to terms that detect binary protein-protein interactions and ‘non-binary’ corresponds to terms that detect potentially indirect associations. An example term in the “invalid” category is “colocalization”. All protein pairs annotated with “invalid” terms were excluded. ‘Binary’ versus ‘non-binary’ evidence was used to categorize protein pairs in the literature-curated dataset as follows. Pairs with no binary experimental evidence were classified as “Lit-NB”, corresponding to 100,940 pairs. Pairs with a single piece of binary evidence and no other evidence were classified as “Lit-BS”, corresponding to 14,477 pairs. Finally, pairs with two or more pieces of evidence including at least one binary evidence were classified as “Lit-BM”, corresponding to 5,589 pairs.

Previous literature-curated datasets generated in 2017 and 2013 were used as a source dataset for pairs experimentally tested in GPCA, MAPPIT and Y2H-v4 (see Engineering of new Y2H destination vectors) experiments. These were generated and processed as above with small differences. Lit-BM-17 and Lit-BS-17 were obtained via the mentha resource data file dated August 28^th^ 2017^30^. Lit-BM-13/Lit-BS-13/Lit-NB-13 were generated as described previously^39^. Yeast PPIs annotated through December 2013 from six source databases: BIND^81^, BioGRID^11^, DIP^10^, MINT^9^, IntAct^82^ and PDB^83^ were extracted and processed using the same protocol.

##### Direct PPIs with experimental structures

The most definitive proof that a pair of interacting proteins are in physical direct contact is the availability of a three-dimensional (3D) structure of their interface. We used the subset of Interactome3D^29^ restricted to experimental structures, excluding homology models. The dataset from the January 2020 release of Interactome3D, referred to as “I3D-exp-20”, was used for most computational analyses. The dataset from the June 2017 release, “I3D-exp-17”, was experimentally tested in its entirety using Y2H v4 (see Engineering of new Y2H destination vectors). The date assigned to PPIs was obtained from the PDB database taking their earliest release date for all PDB structures from the “complete” Interactome3D dataset.

##### Note on the overlap between I3D-exp-20 and Lit-BM-20 PPIs

There were a surprisingly large number of pairs in I3D-exp-20 and not in Lit-BM-20 (1,015 pairs in the difference of I3D-exp-20 from Lit-BM-20 and 746 pairs in the union, see Figure S1B). These pairs are mostly cryo-EM structures (77% Electron Microscopy in the difference vs 36% in the intersection) of larger complexes (median number of entities per structure of 18 in the difference vs 4 in the intersection). The reason for this is that in the generation of the literature-curated datasets (see section Literature-curated biophysical datasets), we don’t use the structural data for direct contacts, we base the binary vs non-binary distinction on the experimental method used and we classify Cryo-EM as non-binary since we don’t know if the reported pairs are in direct contact or not.

##### Predicted structures

The list of PPIs for AlphaFold+RoseTTAFold was from the excel file captioned “Descriptions of all predicted protein-protein interactions”^12^. Six PPIs with missing gene names were discarded. The predicted structures, which were available for pairs with contact probability ≥ 0.9, were downloaded from https://modelarchive.org/doi/10.5452/ma-bak-cepc.

##### Systematic AP-MS

Sys-NB-06 is made up of Gavin *et al.* 2002^6^, Gavin *et al.* 2006^7^, Krogan *et al.* 2006^5^. We didn’t include Ho *et al.* 2002^46^, since it was generated with a smaller, more focussed selection of bait proteins.

**Methods Table 1.**
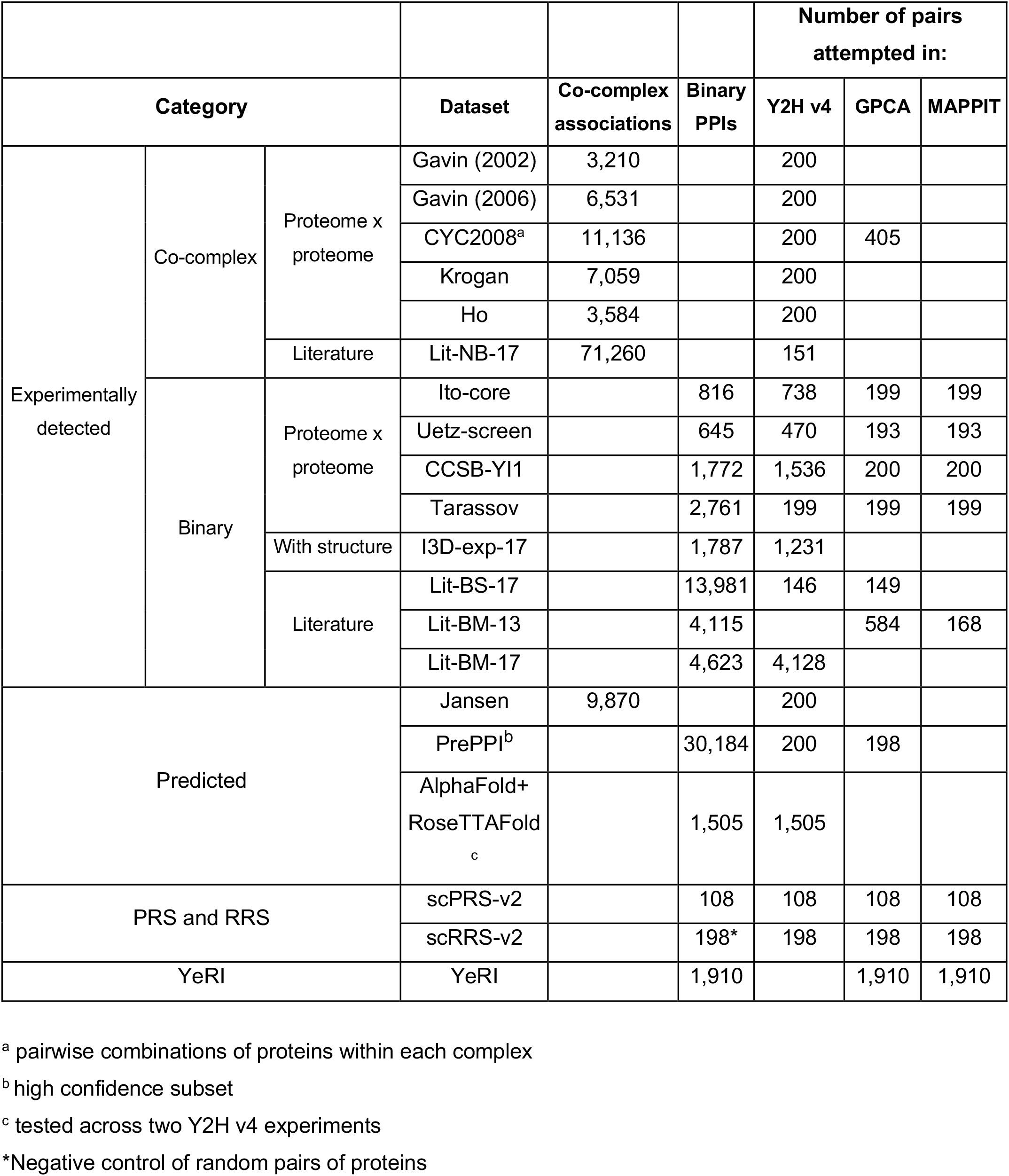
Tested biophysical datasets

##### Functional profile similarity networks (PSNs)

Genetic interaction similarity profile data (GI-PSN) were extracted from Costanzo *et al*. 2016^13^. The average PCC of a pair was used if multiple PCCs were available. Pairs with PCCs ranked in the top 1% were used to generate the GI PSN. Condition-sensitivity data (CS-PSN) was extracted from Hillenmeyer *et al.* 2008^14^. The log of growth ratios from the homozygous deletion data were used to calculate PCC for each pair of genes. Pairs with PCCs ranked in the top 1% were used to generate the condition-sensitivity PSN. Co-expression data (CE-PSN) was downloaded from https://coxpresdb.jp15. The union dataset (Sce-m.c3-0 Sce-r.c1-0, 2018.11.07) was used. Pairs with PCCs ranked in the top 1% were used to generate the co-expression PSN.

**Methods Table 2.** Numbers of genes and interactions in the top 1% percentile of the genetic maps

| PSN | PCC threshold<br>(top 1%) | Number of nodes | Number of edges |
| --- | --- | --- | --- |
| GI | 0.12 | 5,328 | 134,972 |
| CS | 0.42 | 3,479 | 65,147 |
| GE | 0.57 | 3,832 | 99,454 |

#### Generation of scPRS-v2 and scRRS-v2

Due to the change in yeast ORFeome used, we updated our positive reference set (PRS) and random reference set (RRS) from our original set^4^. We named the updated *<u>S</u>accharomyces <u>c</u>erevisiae positive and random* reference sets (scPRS-v2 and scRRS-v2 respectively). In Yu et al., 188 PPIs with five or more papers were finalized as PRS candidates of which 116 had both ORFs in the collection at the time. Of the 188 PPIs, we filtered those pairs to also be in Lit-BM-20, then to have both ORFs in the FLEXGene collection^32^ resulting in a final scPRS-v2 of 108 PPIs. Of 188 RRS pairs in Yu *et al*., we removed all ORFs annotated as dubious, then required they have both ORFs in the FLEXGene collection. To that we increased the size by adding additional pairs randomly selected from the space of all possible pairwise combinations of ORFs in the FLEXGene collection. Since the RRS is used as a negative control, we then filtered out any pairs that appeared in any of the experimental PPI or co-complex association datasets, which resulted in removing one pair that appeared in Lit-NB-20 resulting in a final scRRS-v2 of 198 pairs.

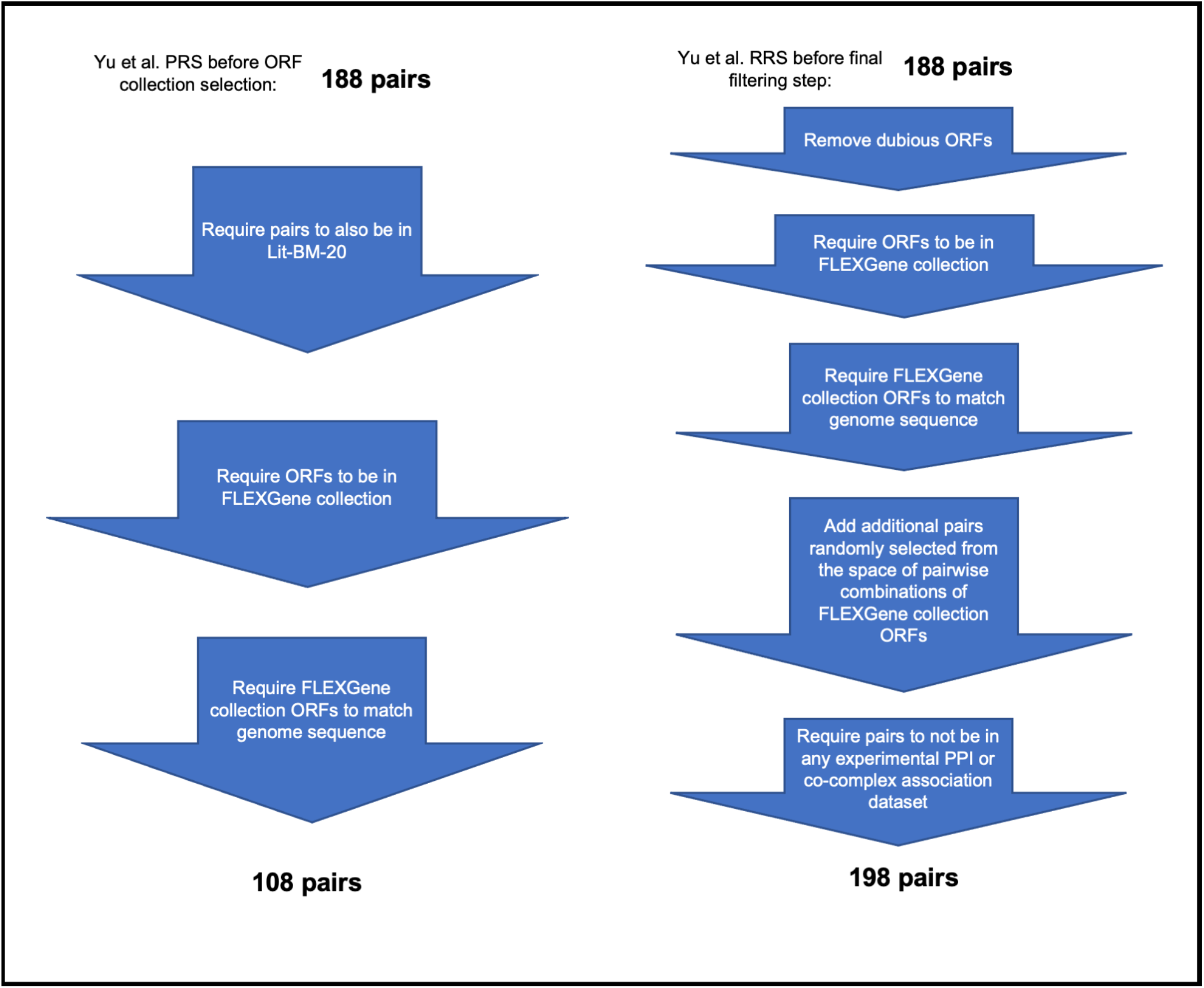

#### Engineering of new Y2H destination vectors

Gateway compatible 2μ high-copy destination vectors pVV212 and pVV213^84^ with a Gal4 DNA binding domain and a Gal4 activation domain, respectively, were modified to be compatible with our standard Y2H vectors pDEST-DB and pDEST-AD-CYH2^72^ with respect to the *LEU2* and *TRP1* as selectable markers. The resulting destination vectors pDEST-DB-QZ212 and pDEST-AD-QZ213 also carry *CAN1* or *CYH2* genes as counterselectable markers, respectively. The *CYH2* and *CAN1* counterselectable markers facilitate plasmid shuffling for the identification of auto-activators^85^. Gateway LR reactions between yeast ORFs flanked by attL1 and attL2 sites with the attR1 and attR2 sites of pDEST-DB-QZ212 and pDEST-AD-QZ213 result in attB1 and attB2 sites flanking yeast ORFs now fused downstream of either the Gal4 DB or Gal4 AD sequences of the respective destination vector. See **Methods Table 3** for detailed information.

**Methods Table 3.** Yeast destination vectors

| **Name** | **pDEST-DB** | **pDEST-DB**<br>**-QZ212** | **pDEST-AD**<br>**-CYH2** | **pDEST-AD**<br>**-QZ213** |
| --- | --- | --- | --- | --- |
| **Fusion partner**<br>**(aa)** | Gal4-DB<br>( 1-147) | Gal4-DB<br>(1-147) | Gal4-AD<br>(768-881) | Gal4-AD<br>(768-881) |
| **Fusion location** | N-terminus | N-terminus | N-terminus | N-terminus |
| **Yeast Promoter**<br>**(nt)** | Truncated *ADH1*<br>(-701 to +1) | Truncated *ADH1* (-<br>410 to +1) | Truncated *ADH1*<br>(-701 to +1) | Truncated *ADH1*<br>(-410 to +1) |
| **Yeast replication**<br>**of origin** | CEN | 2μ | CEN | 2μ |
| **Linker sequence**<br>**between**<br>**3' of Gal4**<br>**element and**<br>**Gateway cloning**<br>**site**<br>**(aa)** | SRSNQ | PEFPS | GGSNQ | ICMAYPYDVPDY<br>ASLGGHMAMEA<br>PS |
| **Yeast terminator** | *ADH1* Term | *ADH1* Term | *ADH1* Term | *ADH1* Term |
| ***E. coli* selection**<br>**marker** | Ampicillin | Ampicillin | Ampicillin | Ampicillin |
| **Yeast**<br>**auxotrophic**<br>**selection marker** | Leucine | Leucine | Tryptophan | Tryptophan |

#### Benchmarking Yeast Two-Hybrid (Y2H) assay versions

Assay versions were benchmarked using scPRS-v2 and scRRS-v2. The new Y2H version with destination clones in vectors pDEST-DB-QZ212 and pDEST-AD-QZ213 was named Y2H version 4 (Y2H v4). Y2H v1 - v3 can be found in Luck et al, Nature, 2020^20^. The performance of Y2H v4 was compared to Y2H v1, which consists of destination clones in pDEST-AD-*CYH2* and pDEST-DB, and was used to generate CCSB-YI1^4^. The Y2H assay was performed as described previously^39, 72^. Briefly, Y8930:pDEST-DB-QZ212-ORF and Y8800:pDEST-AD-QZ213-ORF haploid strains were inoculated and mated. After enrichment for diploids in SC-Leu-Trp, diploids were spotted on SC-Leu-Trp-His+3AT solid media, testing for *GAL1::HIS3* activation and on a set of SC-Leu-His+3AT plates supplemented with 10 mg/L cycloheximide (CHX) to identify spontaneous DB-ORF auto-activators^72^. After 3 days incubation at 30°C, yeast strains growing on SC-Leu-Trp-His+3AT solid media and not on SC-Leu-His+3AT+CHX media were scored as positives. The interacting pairs were identified based on plate position.

#### Generation of an expanded yeast ORFeome collection

Yeast FLEXGene clone collection^32^ of full length ORFs cloned in either pDONR201 or pDONR221, both Kan^R^, contains 4,933 ORFs, after removal of redundant ORFs and ORFs that no longer match SGD-annotated ORFs (version 2014) (https://www.yeastgenome.org/). For the remaining 950 SGD-annotated ORFs not in Yeast FLEXGene, entry clones were generated in-house and are referred to as supplemental ORFeome collection. ORF sequences were amplified without their native stop codon sequences from either *S. cerevisiae* S288C genomic DNA (ORFs without introns) or cDNA (ORFs containing introns) using KOD high fidelity polymerase (Novagen) and 18-20 nucleotide ORF-specific forward and reverse PCR primers tailed with Gateway attB1 and attB2 sequences attB1 Forward primer tail 5’ GGGGACAAGTTTGTACAAAAAAGCAGGCTCCACC attB2 Reverse primer tail 5’ GGGGACCACTTTGTACAAGAAAGCTGGGTCCTA from Hu et a^32^, respectively, essentially as described^86^. The CTA sequence in the Gateway tail of the reverse primer provided a synthetic stop codon for all ORFs. Amplified ORFs were transferred to pDONR223 (Spec^R^) by Gateway BP recombination cloning (Invitrogen) and transformed into chemically competent DH5α *E. coli* cells. Sanger sequencing of PCR products, generated with universal forward and reverse primers, was used to confirm the identity of all cloned ORFs as described^86^. 921 ORFs were obtained using this approach.

#### ORFeome cloning in Y2H destination vectors

To generate an arrayed library of DB-ORF and AD-ORF hybrid proteins, the yeast ORFs were transferred into both destination vectors, pDEST-DB-QZ212 and pDEST-AD-QZ213, by Gateway LR recombination cloning (Invitrogen). Gateway LR reaction products were transformed into DH5α *E. coli* cells, plasmid DNA was extracted and used to transform yeast strains. pDEST-DB-QZ212 and pDEST-AD-QZ213 expression clones were transformed into yeast strains *MAT*α Y8930 and *MAT***a** Y8800, respectively^72^.

#### Auto-activator detection for filtering before Y2H screening

We tested for auto-activation of the *GAL1::HIS3* reporter gene by AD-ORF or DB-ORF fusion proteins in both haploid and diploid yeast cells. To identify auto-activator clones in haploid yeast, Y8930:DB-ORF and Y8800:AD-ORF strains were grown to saturation in SC medium lacking Leucine (SC-Leu) or Tryptophan (SC-Trp), respectively. After 24 hours of incubation, Y8930:DB-ORF and Y8800:AD-ORF haploids were spotted on SC-Leu-His+3AT or SC-Trp-His+3AT to test for *GAL1::HIS3* activation. Viability of the haploids was confirmed with growth on SC-Leu or SC-Trp, respectively.

To identify auto-activators in diploid yeast, *MAT*α Y8930:DB-ORF and *MAT***a** Y8800:AD-ORF strains were mated against their respective opposite mating type strains carrying the corresponding destination vectors without any fused ORFs. Mating was conducted in rich medium, YEPD, and resulting diploids were enriched following growth in SC-Leu-Trp. Diploids were spotted on SC-Leu-Trp-His+3AT, to test for *GAL1::HIS3* activation, and on SC-Leu-Trp to confirm the viability of the diploids. For both haploids and diploids, after incubation at 30°C for 3-4 days, strains growing in the absence of histidine were considered auto-activators. 560 DB-ORFs and 1 AD-ORF were removed from the final screening collection.

The remaining DB-ORF and AD-ORF clones were re-arrayed into four different groups to separate ORFs with similar nucleotide sequences, defined as BLAST scores of x100 and above. Separation of similar ORFs makes the downstream sequence identification of the short NGS reads more accurate, as the reads are aligned to specific groups of ORFs without sequence ambiguity. Filtering for pairs that passed autoactivator screening and successful cloning resulted in a final collection which was then used for systematic screening included 4,778 DB-ORF clones and 5,700 AD-ORF clones, covering a total of 5,854 yeast ORFs.

#### Primary yeast two-hybrid (Y2H) screening

Three replicate Y2H screens were performed. Individual *MAT*α Y8930:DB-ORFs were mated in YEPD against a pool of ∼700 (FLEXGene collection) or ∼200 (supplemental collection) *MAT***a** Y8800:AD ORFs. AD-ORF pool size was decreased for the supplemental collection to facilitate screening. After enrichment in SC-Leu-Trp, 5µl of the culture was spotted on SC-Leu-Trp-His+3AT solid media and on SC-Leu-His+3AT+ 10mg/L CHX to identify spontaneous DB-ORF auto-activators^72^. After incubation at 30°C for 3 days, strains growing on SC-Leu-Trp-His+3AT but not on SC-Leu-His+3AT+CHX were picked and grown in liquid SC-Leu-Trp. As we used libraries of pools of *MAT***a** Y8800:AD-ORF, it is possible to obtain more than one interaction per mini-library. To account for that, we picked up to three colonies per growth spot. Cell lysates were prepared from the saturated cultures and used as templates in PCR reactions to amplify and identify the bait and prey sequences^72^.

#### Yeast colony sequencing

To efficiently and cost-effectively identify both bait and prey proteins for thousands of positive colonies, we used a method called SWIM-seq (Shared-Well Interaction Mapping by sequencing) as described^20^. Briefly, DB and AD-ORFs were simultaneously amplified from 3μl yeast lysate, using well-specific primers. PCR reactions were performed using Platinum Taq (Life Technologies). After PCR amplification, barcoded PCR products from an entire 96 well plate were pooled together and purified (Qiagen, PCR Purification Kit). These pooled amplicons from each plate were subjected to Nextera “tagmentation” using Tn5 transposase to generate a library of amplicons with random breaks to which the adapters have been ligated^87^. We then re-amplified those fragments to generate a library of amplicons such that one end of each amplicon bears the well-specific tag and the other “ladder” end bears the Nextera adapter. A final Illumina sequencing library was prepared by adding plate indexes using the i5 and i7 Illumina adapter sequences. Next generation sequencing was performed with Illumina Solexa technology allowing for identification of interacting first pass pairs of proteins (FiPPs) (see Sequence identification of interacting ORFs). Due to the small number of pairs to be identified, interacting pairs from the first screen of the supplemental space were amplified with the universal AD and DB forward and reverse primers and ORF sequences were identified by Sanger sequencing (Genewiz). All SWIM-primers (**Methods Table 4**) were synthesized by Thermo Fisher Scientific, whereas the universal AD, DB and term primers were synthesized by Eurofins Genomics.

**Methods Table 4.**
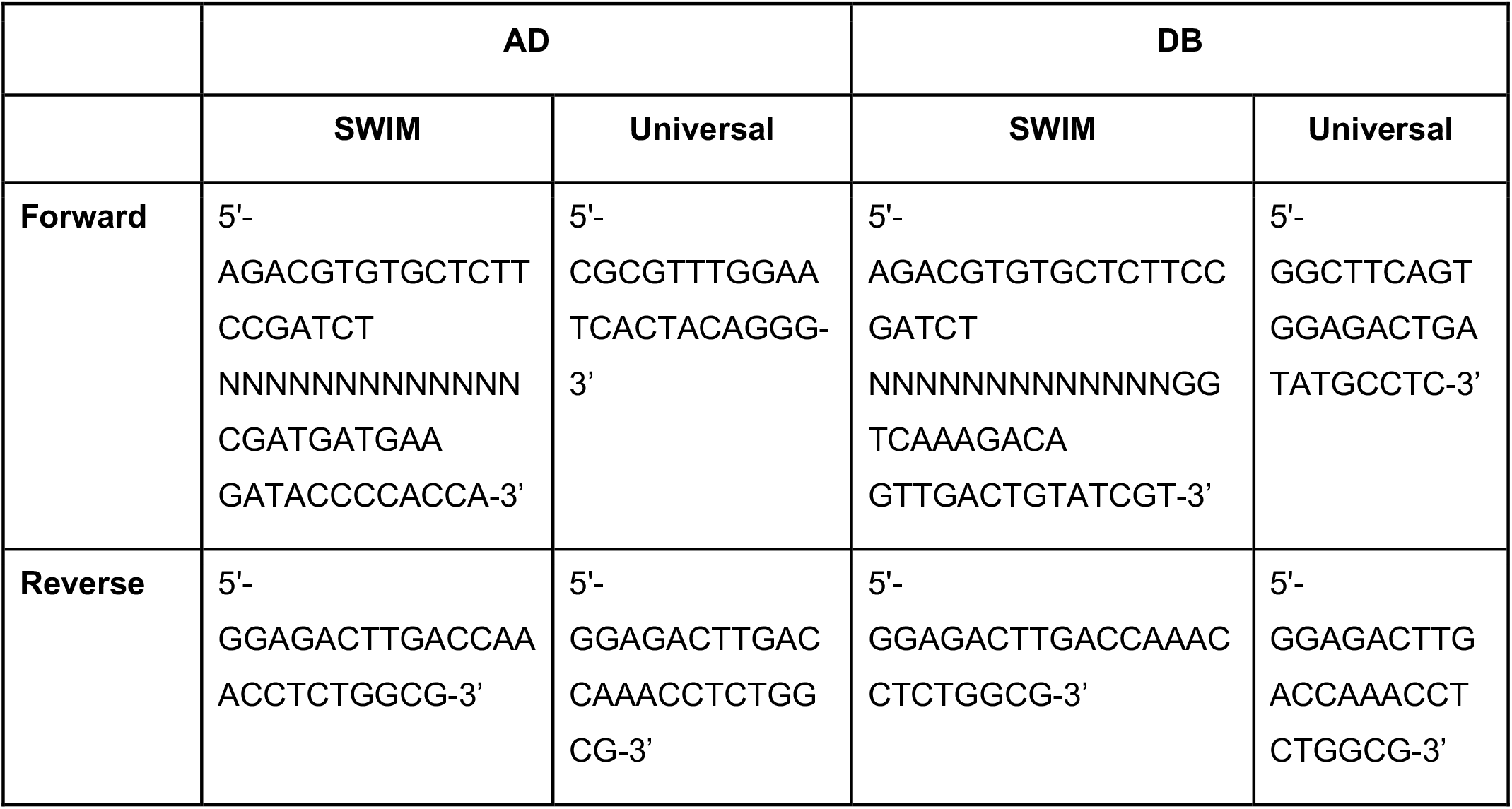
Primers used (Ns denotes 13-mer well index)

#### Pairwise test

To confirm all FiPPs, a pairwise test was performed in the same DB-X/AD-Y orientation they were found in the primary screens. Briefly, glycerol stocks from Y8930:DB-ORF and Y8800:AD-ORF haploid strains were inoculated in SC-Leu or SC-Trp, respectively. Saturated cultures were mated in YEPD. After enrichment for diploids, yeast were spotted on SC-Leu-Trp-His+1 mM 3AT solid media, testing for *GAL1::HIS3* activation. Preliminary investigations using four technical replicates demonstrated that in 97% of the cases, the quadruplicates behaved identically (data not shown). Therefore, given the high reproducibility of technical replicates, the culture was spotted only once per selective media. To increase the robustness of our approach we implemented an additional test to identify *de novo* auto-activators in which Y8930:DB-ORF strains were mated against a Y8800:AD with no ORF fused to the activation domain (Y8800:AD-Empty ORF) and spotted on SC-Leu-Trp-His+1 mM 3AT solid media. Diploids that gave rise to growth on SC-Leu-Trp-His+1 mM 3AT media, but did not grow when the respective Y8930:DB-ORF was mated to Y8800:AD-Empty ORF, were selected as positive interacting pairs of proteins. Positive protein pairs were sequence confirmed as done for the primary screens as described above. As positive and negative controls, the scPRS-v2 and scRRS-v2 pairs were distributed randomly across the respective mating plates and tested at the same time. For a batch of pairwise testing to be considered successful we required no more than 1% of RRS and between 10-25% of PRS to be scored positive.

#### Validation in orthogonal assays

To assess the precision of various datasets^88^, PPIs were validated in two orthogonal assays: <u>Ma</u>mmalian <u>p</u>rotein-<u>p</u>rotein interaction trap (MAPPIT)^33^ and *<u>G</u>aussia princeps* luciferase <u>p</u>rotein <u>c</u>omplementation <u>a</u>ssay (GPCA)^34^. As positive and negative controls, we used pairs of scPRS-v2 and scRRS-v2 respectively. For both assays, expression clones were generated by Gateway LR recombination cloning as described above. Expression clones for GPCA were generated by transferring ORFs into pSPICA-N1 and pSPICA-N2 destination vectors^34^, each expressing a different fragment of humanized *Gaussia princeps* luciferase (GL1 and GL2)^89^. MAPPIT expression clones were generated by LR transfer of ORFs into pMBU-I-2994 and pMBU-I-4199 destination vectors^33^. After transformation of all expression clones into DH5α *E. coli* cells, plasmid DNA was extracted and purified using Qiagen 96 Turbo kits (Qiagen) on a BioRobot 8000 (Qiagen). Three different GPCA and two different MAPPIT experiments were performed.

##### GPCA

GPCA experiments were performed as described previously^34^. Briefly, on the first day of the assay, ∼30,000 to 40,000 HEK293T cells were seeded in each well of a 96 well microtiter plate (Greiner Bio-One). DNA concentration was measured for all clones and samples were diluted to a final concentration of 25ng/μl. After a 24-hour incubation at 37°C, confluent cells were transfected with 300ng of pSPICA-N1-ORF and pSPICA-N2-ORF vectors using polyethylenimine (PEI). After a second 24-hour incubation at 37°C, cells were washed with PBS supplemented with calcium and magnesium chloride. To lyse the cells 40μl of 5x diluted Renilla lysis buffer (Promega) were added to each well. The plate was then covered with aluminum foil and agitated at 900 rpm for 30 minutes at 37°C for cell lysis. Luciferase activity was measured on a TriStar Berthold Microplate reader by adding 50μl per well of Renilla luciferase substrate (Renilla Luciferase Assay System, Promega), with a measurement time of 4 seconds. The measurement score, RLU (relative light unit), was assigned to the tested pair.

##### MAPPIT

As an orthogonal validation assay, MAPPIT experiments were performed as described elsewhere^20, 39^. In short, HEK293T cells were grown in 384-well plates and co-transfected with a luciferase reporter and plasmids for both bait and prey fusion proteins. Twenty-four hours post-transfection, cells were either stimulated with ligand (erythropoietin) or left untreated, then incubated for an additional 24 hours before luciferase activity was measured in duplicate. The MAPPIT validation experiment was deemed valid, if both bait and prey were successfully cloned into expression vectors and bait expression was detected using a chemiluminescence meter. “Fold-induction” values (signal from stimulated cells divided by signal from unstimulated cells) were calculated for each tested pair, and two negative controls (no bait with prey and bait with no prey). Each tested pair was assigned a quantitative score: the fold-induction value of the pair divided by the maximum of the fold-induction value of the two negative controls.

#### Experimental benchmarking of public PPI datasets

PPIs extracted from the biophysical maps described in **Methods Table 1** have been tested in assays Y2H v4, GPCA and MAPPIT following the same experimental procedures as described above. A summary of the number of tested pairs in each dataset is available in **Methods Table 1**. Samples, if used, were drawn randomly.

An additional Y2H v4 experiment was performed to test pairs from the AlphaFold+RoseTTAFold dataset, along with scPRS-v2 and scRRS-v2. Roughly half of AlphaFold+RoseTTAFold PPIs had already been tested in the first Y2H v4 experiment, as they overlapped with one or more of the tested datasets. So, in the additional Y2H v4 experiment we only tested pairs that had not been tested in the first experiment. After checking that the scPRS-v2 and scRRS-v2 results were consistent between the two experiments, the overall recovery of AlphaFold+RoseTTAFold was calculated combining the data from both experiments, with every pair having been tested in exactly one of the two experiments.

#### Direct or indirect contact in a complex structure

We queried Interactome3D (version 2020_01)^29^ for complexes involving three or more proteins with an experimental structure available. For all combinations of protein pairs within a complex, Interactome3D calculated the number of residue-residue contacts by accounting for hydrogen bonds, van der Waals interactions, and salt and disulfide bridges. We defined protein pairs with five or more contacts as direct, and remaining pairs as indirect. Using this annotation for each dataset, the fraction of direct PPIs was calculated as the number of direct PPIs reported in the dataset divided by the number of direct and indirect pairs reported in the dataset.

#### K_d_ dataset

Yeast PPIs with measured dissociation constant (*K_d_*) values were obtained from the PDBbind database^90^ 2017 release and from^91^. In the case where multiple values existed for a pair, the geometric mean was used.

#### PPIs in KEGG pathways and in the four gold standard inner- and outer-complexome datasets

We collected PPIs from KEGG annotated as activation, inhibition, phosphorylation, dephosphorylation, ubiquitination, glycosylation, methylation, binding/association, complex as defined by KEGG. Gene expression relations and enzyme-enzyme relations were excluded. The four gold standard inner- and outer-complexome PPI datasets are: i) direct co-complex PPIs using the intersection between protein complex dataset collected by Costanzo *et al.* 2016 filtered with three or more subunits and direct interactions from Interactome3D (Direct co-complex); ii) co-complex pairs annotated in 5 KEGG yeast pathways Cell Cycle, Meiosis, MAPK Signaling pathway, Autophagy and Mitophagy (KEGG co-complex); iii) PPIs regulating activation or inhibition from the same 5 KEGG yeast pathways (KEGG regulation); and iv) high-quality kinase-substrate pairs from the Yeast KID database (http://www.moseslab.csb.utoronto.ca/KID/)^50^ with score greater or equal to 6.4 (p-value < 0.01) (Kinase-substrate).

#### List of genes of unknown function

A list of 979 *S. cerevisiae* genes of unknown function was obtained from Table S9 of Wood et al. 2019^63^, of which 950 were within the list of yeast ORFs considered for this study (see section Yeast protein-coding ORFs).

#### Protein properties

1. Number of publications per gene was extracted from the gene2pubmed file from NCBI, downloaded on 2018-08-01.
2. Protein abundance information was downloaded from PaxDB (https://pax-db.org) undetected pairs were given an abundance of 0.
3. Gene essentiality information was downloaded from the *Saccharomyces* Genome Deletion Project (https://www.yeastgenome.org).
4. Conservation score was derived by combining data from HomoloGene (ftp://ftp.ncbi.nih.gov/pub/HomoloGene/build68) and Carvunis *et al.* 2012^92^. For a gene with homologs in HomoloGene, its conservation score is the number of distinct non-*S. cerevisiae* species that it shares the same homologene group with plus 9, assuming that it is conserved in the 10 Ascomycota species. For genes without homologs in HomoloGene, we used classification proposed in Carvunis *et al* where genes were scored from 1-10 based on their conservation throughout the Ascomycota phylogeny. Genes without homologs in HomoloGene and that did not appear in the Carvunis data were given a score of 0.
5. Complex size was the number of different protein subunits taken from the complexome dataset. If a protein was a member of multiple complexes, the size of the largest complex was used.

#### Treatment of heterodimers and homodimers

Unless otherwise noted, homodimers were excluded from most analyses since comparisons between physical interactions and functional relationships are obviously not applicable to single genes (all PCC values of functional profiles would be 1.0 by definition).

#### Calculation of recovery rates in Y2H v4, MAPPIT and GPCA

In MAPPIT and GPCA assays, pairs were scored positive or negative based on thresholds set by the highest scoring scRRS-v2 pair in the corresponding experiment. For all three assays, pairs without valid quantitative scores were dropped, and recovery rates were calculated as the number of positive pairs over the sum of the positive and negative pairs. The error bars on the recovery rates were calculated using a Bayesian model (a binomial likelihood with a uniform prior), taking the central 68.27% interval of *Beta* (*p* + 1, *n* + 1), where *p* and *n* are the number of pairs testing positive and negative, respectively. P-values for difference in recovery between two datasets tested in the same experiment are calculated using Fisher’s exact test, two-sided in all cases except when testing a dataset against the scPRS-v2 / scRRS-v2 positive or negative controls, where a one-sided test is used. For the AlphaFold+RoseTTAFold Y2H v4 results, where the data was split across two experiments, the scPRS-v2 recovery is calculated as an average of the two experiments, weighting by the number of positive AlphaFold+RoseTTAFold pairs in each experiment.

#### Calculation of interface areas of PPIs

We retrieved experimental structures using Interactome3D version 2018_04^29^. For each subunit in a complex structure, we defined its interaction interface as the residues for which the Accessible Surface Area (ASA) changed more than 1 Å^2^ between the bound and unbound state.

#### Prediction of ΔG

We used FoldX 5^93^, first running RepairPDB, then Optimize, then AnalyseComplex, all with default parameters.

#### Interaction 2D histogram heat maps

For a particular gene/protein property and a network, we ranked all proteins using that property. Tied values were sorted randomly. The proteins were split into an equal number of bins, creating 2D bins of the protein-by-protein space. Number of edges in the diagonal bins were multiplied by a factor of *N^2^ / (N^2^ / 2 - N / 2)*, where *N* is the number of proteins in the bin, to correct for the smaller number of possible pairwise combinations, since edges were undirected. Homodimeric interactions were excluded. In the case where we corrected the CS-PSN heatmaps for the untested essential genes, we divided the count in each bin by the fraction of pairs where both genes were tested in generating the CS-PSN data.

To calculate the p-values for each 2D bin, we randomly shuffled the order of the proteins 1,000 times. In each permutation of the proteome we calculated the 2D histogram counts, recorded the maximum and minimum bin count (to account for the multiple testing effect of having many bins) and calculated the p-value, for each bin, as the fraction of the random maximum/minimum counts that the observed count is above/below, multiplied by two to account for the two-tailed nature of the test. This was done separately for diagonal and off-diagonal bins because there are a different number of possible combinations of undirected edges between them.

#### Sequence identification of interacting ORFs

We used an existing computational pipeline^20^ to process demultiplexed paired-end reads returned from Illumina sequencing and identify the interacting ORF pairs from the Y2H screen. Paired-end reads are in fastq format, with one read, R1, containing a part of the ORF sequence and the other paired read, R2, containing the well index. We used Bowtie 2^16^ (v2.2.3) to align all R1 reads to reference sequences and extracted the well-identifying indices from the R2 reads. AD-ORFs and DB-ORFs that shared the same well indices were paired together and called FiPPS. To identify likely true AD/DB pairs, we developed a “SWIM score”^20^ *S* that takes into account the AD and DB reads in each well, total reads returned from the sequencing run, and other factors.

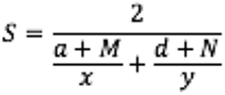

where *x* and *y* are read counts of an AD-ORF and DB-ORF in a given well respectively, *a* and *d* are total read counts of all aligned AD-ORF and DB-ORF in that well, and *M* and *N* are pseudo-counts for AD and DB respectively, which were constant for each sequencing batch but varied for different batches. We then selected FiPPs for pairwise testing using a cutoff that balances the risk of testing too many false positives FiPPs versus not testing too many true positive FiPPs. The cutoff varied for different screens and sequencing runs to adjust for slight variations in the screening and sequencing protocol.

#### Calculation of enrichment for connecting proteins in the same subcellular compartment, pathway, and complex

Subcellular compartment data was^65^ obtained from CYCLoPS^65^, using the WT data, annotating a protein to a compartment if it has any non-zero value in any of the three repeats. Pathways were obtained from KEGG^49^. Complexes were obtained from CYC2008^45^. The number of PPIs that connected two different proteins in the same compartment, pathway or complex was divided by the mean value for 1,000 degree-preserved randomized networks, generated using the Viger and Latapy algorithm implementation through python iGraph^94^, and CI values were taken from the innermost 68.27% of the random networks.

#### SAFE network visualization

We used the SAFE network visualization tool (v1.5)^35^. The layouts were generated with Cytoscape (v3.4.0)^95^ using the edge-weighted spring embedded layout. GO terms were downloaded from SGD database (version on Jan 17^th^ 2019) and GO^35^ terms enriched with P < 0.05 were colored and labeled. SAFE analysis was run with the default option except layoutAlgorithm = none (using layout generated by Cytoscape), neighborhoodRadius = 200, and neighborhoodRadiusType = absolute.

#### Estimates of the complete yeast interactome size

We used three estimates, relying on partially overlapping assumptions and data, made by independent groups, that predicted the yeast protein binary interactome contains between ∼18,000 and ∼38,000 direct binary interactions, corresponding to ∼0.1-0.2% of all ∼18,000,000 possible protein pair combinations^4^.

- From Yu *et al.* 2008^4^ 18,000 (13,500-22,500 95% CI). Taken from Page 107: “we estimated that the yeast binary interactome consists of ∼18,000 +-4500 interactions (SOM VI)” From SOM VI the +/-refers to the 95% CI.

- From Stumpf *et al.* 2008^36^ 28,472 (26,650-30,460 95% CI). Taken from the Uetz et al. numbers from Table 1. We use the estimate made using Uetz et al. because three of the other datasets contain indirect protein-protein associations (Ho et al., Gavin et al. and DIP) and the estimate using Ito et al. uses the full dataset, mainly made up of the ‘Ito-noncore’ subset that was shown to be of poor quality when retested Y2H and PCA^4^.
- From Sambourg and Thierry-Mieg 2010^37^ 37,600 (32,252-43,472 95% CI). Taken from Page 6: “Taken together, this allows to estimate the size of the binary yeast interactome at ∼ 37,600 interactions (95% confidence interval: 32252-43472, constructed with the normal approximation method).”

One relatively minor difference between the estimates is that Stumpf et al. are considering only heterodimeric PPIs whereas Yu et al. and Sambourg et al. are also counting homodimeric PPIs and so we account for this when estimating the fraction of predicted interactome mapped by excluding homodimers for the Stumpf et al. estimate and including them for the Yu et al. and Sambourg et al. estimates.

#### Prediction of gene functions using guilt-by-association approach

In the guilt-by-association approach the function of a node is inferred from the function of its neighbors. In particular, for each node we count the number of its neighbors annotated with a given function (*n*). This score is then compared to a random benchmark, obtained by randomizing the network 10,000 times in a degree-preserved way. Calculating the *z*- score, *z* = (*n* – 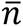) ÷ α, is the traditional way of such comparison, obtained by standardizing the original score with the expectation value (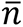) and standard deviation (σ) of the score that would be expected by chance. Yet, the *z*-score is not free from degree biases and prefers low degree nodes with extremely small σ. We therefore apply a related measure, called the effect size. The effect size *n* − (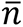 + -+*ασ*)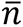 is obtained by comparing the original score with the reasonably expected value of the random benchmark, estimated as the mean value (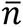) and α-times the standard deviation (σ). In practice, we use α = 2, selecting the same candidates as a traditional z-score threshold of *z* ≥ 2, but ordering them based on the amount of signal beyond random expectations to avoid a bias towards low-degree nodes. Functional annotations of genes with GO Biological Process terms were obtained as described above and further restricted to annotations with the experimental evidence codes EXP, IDA, IPI, IMP, IGI, IEP, HTP, HDA, HMP, HGI, and HEP.

#### Network fragmentation

The error bands are generated from randomly sampling a fraction of the PPIs from each network, where the fraction varies from 5% to 95% in 5% increments, with 1,000 random subnetworks generated at each point.

#### Protein complex subnetworks

For each protein complex, direct interactions were defined by I3D-exp-20, described above, indirect associations were all protein-protein combinations where both proteins appeared in the same experimental structure but not in direct contact.

#### General protein categories

The categorisation of proteins into “Genetic information processing”, “Metabolism”, and “Not mapped” was obtained from level 1 in the KEGG-based mapping of Liebermeister *et al.* 2014^62^.

#### Degree distribution plots

Degree distributions were plotted according to Chapter 4, Advanced Topic 3.B of Network Science^96^, on a log-log scale with logarithmic binning, with the unbinned data shown in grey.

#### Overlap calculation between biophysical and functional networks

For each biophysical network and several KEGG pathways, we measured the fraction of interactions that are also connected in each of the functional networks defined above, discarding homodimeric PPIs. We calculated the overlap by dividing the number of interactions in the PPI network also found in the functional network by the total number of interactions in the PPI network where both proteins were present in the search space of the functional network. The error bars were calculated using a Bayesian model (a binomial likelihood with a uniform prior), taking the central 68.27% interval of *Beta* (*p* + 1, *n* + 1), where *p* and *n* are the number of pairs testing positive and negative, respectively.

#### Date and party hubs

Co-expression data was obtained from COXPRESdb^15^. To ensure robustness against the exact definition of date and party hubs, three different cutoffs were used, hubs were defined as proteins with a degree in the top 5% or 10% in each network, or those with degree ≥ 10. PCC cutoffs of 0.3 and 0.35 were used, where proteins with a mean coexpression PCC across all partners above the cutoff were party hubs and below the cutoff were date hubs.

#### Overlap by degree plots

For each combination of a biophysical and functional network, we conducted a logistic regression, on the dataset of biophysical interactions, where the binary dependent variable represents whether or not the two proteins of the biophysical interaction are also connected by an edge in the functional network, and the single independent variable is the higher of the two degrees, in the biophysical network, of the interacting proteins. The max degree per PPI variable is log2 transformed. Only PPIs where the pair of proteins were tested in generating the functional network were used. Shaded error bands represent 95% CI. Binned data is also shown, with 10 evenly sized bins, with the binned data displayed on the x-axis at the mean max degree value of the bin.

## Description of Supplementary Tables

**Supplementary Table 1.** The literature curated PPI datasets used in this study from 2020.

**Supplementary Table 2.** The curated evidence for literature PPI datasets in 2020.

**Supplementary Table 3.** The literature curated PPI datasets used in this study from 2013.

**Supplementary Table 4.** The literature curated PPI datasets used in this study from 2017.

**Supplementary Table 5.** Various biophysical and functional networks used in this study.

**Supplementary Table 6.** A list of the ORFs screened in generating YeRI.

**Supplementary Table 7.** Results from the experiment to compare Y2H assay versions.

**Supplementary Table 8.** scPRS-v2 the positive reference set of PPIs.

**Supplementary Table 9.** scRRS-v2 the reference set of random pairs.

**Supplementary Table 10.** Results from experiments performed with the GPCA and MAPPIT PPI assays.

**Supplementary Table 11.** The yeast reference interactome (YeRI) PPI dataset.

**Supplementary Table 12.** ABBI-21 (Atlas of Binary Biophysical Interactions), which is the union of Uetz-screen, Ito-core, CCSB-YI1 and YeRI.

**Supplementary Table 13.** Predicted GO terms for genes of unknown function using a guilt-by-association approach based on the GO terms of their interaction partners in ABBI-21 and YeRI.

**Supplementary Table 14.** Results from the Y2H v4 pairwise test.

**Supplementary Table 15.** Results from an additional Y2H v4 pairwise test of the remaining AlphaFold+RoseTTAFold PPIs.

**Supplementary Table 16.** Various protein properties for each ORF considered in this study.

**Supplementary Table 17.** Pairs of proteins within a complex and whether they are in indirect or direct contact. Data derived from experimental structures.

**Supplementary Table 18.** Interface area and predicted *ΔG* for experimental structures from I3D-exp-20.

**Supplementary Table 19.** Interface area and predicted *ΔG* for predicted structures from AlphaFold+RoseTTAFold.

